# Local Shape Descriptors for Neuron Segmentation

**DOI:** 10.1101/2021.01.18.427039

**Authors:** Arlo Sheridan, Tri Nguyen, Diptodip Deb, Wei-Chung Allen Lee, Stephan Saalfeld, Srini Turaga, Uri Manor, Jan Funke

## Abstract

We present a simple, yet effective, auxiliary learning task for the problem of neuron segmentation in electron microscopy volumes. The auxiliary task consists of the prediction of Local Shape Descriptors (**LSDs**), which we combine with conventional voxel-wise direct neighbor affinities for neuron boundary detection. The shape descriptors are designed to capture local statistics about the neuron to be segmented, such as diameter, elongation, and direction. On a large study comparing several existing methods across various specimen, imaging techniques, and resolutions, we find that auxiliary learning of LSDs consistently increases segmentation accuracy of affinity-based methods over a range of metrics. Furthermore, the addition of LSDs promotes affinity-based segmentation methods to be on par with the current state of the art for neuron segmentation (Flood-Filling Networks, FFN), while being two orders of magnitudes more efficient—a critical requirement for the processing of future petabyte-sized datasets. Implementations of the new auxiliary learning task, network architectures, training, prediction, and evaluation code, as well as the datasets used in this study are publicly available as a benchmark for future method contributions.

## 1 Introduction

The goal of connectomics is the reconstruction and interpretation of neural circuits at synaptic resolution. These wiring diagrams provide insight into the inner mechanisms underlying behavior and help drive future theoretical experiments (Schneider-Mizell et al., 2016; Motta et al., 2019; Bates et al., 2020; Hulse et al., 2020). Additionally, the generation of connectomes has proven to complement existing techniques such as calcium imaging and electrophysiology where the resolution is often not sufficient to parse the circuitry in detail (Schlegel et al., 2016; Turner-Evans et al., 2020).

Currently, only electron microscopy (EM) allows imaging of neural tissue at a resolution sufficient to resolve individual synapses and fine neural processes. Two popular methods for imaging these volumes are serial block-face scanning EM (SBF-SEM) and focused ion beam scanning EM (FIB-SEM). While the former technique is faster and generates high lateral resolution, it results in lower axial resolution due to section slicing. The latter method produces isotropic resolution by etching the face of the volume with a focused ion beam before imaging. However, this method is slower than serial section approaches. Briggman and Bock (2012) provide a thorough overview of these imaging approaches and others, including ssTEM and ATUM-SEM. All methods have been used to generate invaluable datasets for the connectomics community (Lee et al., 2016; Zheng et al., 2018; Dorkenwald et al., 2019; Schneider-Mizell et al., 2020; Turner et al., 2020; Scheffer et al., 2020; Phelps et al., 2021).

Depending on the specimen and the circuit of interest, current EM acquisitions produce datasets ranging from several hundred terabytes to petabytes. For instance, the raw data of a full adult fruit fly brain (Fafb) comprises ∼50 teravoxels of neuropil (Heinrich et al., 2018). Even sub-volumes taken from vertebrate brains, which do not contain brain-spanning circuits, result in massive amounts of data: The authors of Kornfeld et al. (2017) imaged a region from a zebrafinch brain containing ∼10^6^ µm^3^ (∼663 gigavoxels) of raw data. More recently, a larger volume of mouse visual cortex has been imaged, comprising ∼3 × 10^6^µm^3^ (∼6,614 gigavoxels) (Dorkenwald et al., 2019; Turner et al., 2020; Schneider-Mizell et al., 2020; Yin et al., 2020; Microns Consortium et al., 2021). A 1.4 petabyte volume taken from human cortex further demonstrates the rapid advances in massive dataset acquisition (Shapson-Coe et al., 2021). In order to reconstruct circuits in a full mouse brain, however, the authors of Abbott et al. (2020) argue that it will require an acquisition of around one exabyte of raw data (one million terabytes).

With datasets of this magnitude, purely manual reconstruction of connectomes is infeasible. On average, manual tracing in mouse tissue takes ∼1-2 hours per millimeter (Boergens et al., 2017; Motta et al., 2019). Larval tissue averages ∼13.7 hours per millimeter (Schneider-Mizell et al., 2016), which is comparable to tracing speeds in the *Drosophila* dataset, Fafb (Zheng et al., 2018) due to the challenging nature of invertebrate neuropil. Even the small brain of a *Drosophila* contains an estimated 100,000 neurons, which would require ∼125 years of manual effort to trace each neuron to completion.

Consequently, automatic methods for the reconstruction of neurons and identification of synapses have been developed. Over the past decade, methods targeting relatively small volumes have pioneered the reconstruction of neurons (Turaga et al., 2010; Lee et al., 2017), and synapses (Kreshuk et al., 2015; Buhmann et al., 2018). More recently, these efforts have been improved to tackle the challenges of large datasets for neurons (Januszewski et al., 2018; Funke et al., 2019; Dorkenwald et al., 2019; Li et al., 2019), synaptic clefts (Heinrich et al., 2018), and synaptic partners (Huang et al., 2018; Buhmann et al., 2020). With the help of an automatic neuron segmentation method, neuron tracing times decreased by a factor of 5.4 to 11.6 minutes per µm path length (Li et al., 2019), effectively trading compute time for human tracing time.

However, given the daunting sizes of current and future EM datasets, limits on available compute time become a concern. Future algorithms do not only need to be more accurate to further decrease manual tracing time, but also computationally more efficient to be able to process large datasets in the first place. Consider the computational time required by the current state of the art, FFN: Assuming linear scalability and the availability of 1000 contemporary GPUs (or equivalent hardware), the processing of a complete mouse brain would take about 226 years. This example alone goes to show that the objective for future method development should be the minimization of the total time spent to obtain a connectome, including computation and manual tracing. Therefore, automatic methods for connectomics need to be fast, scalable (*i.e*., trivially parallelizable), and accurate.

### 1.1 Neuron Segmentation Methods

Neuron reconstruction is an instance segmentation problem. Unlike semantic segmentation, in which the goal is to assign every voxel to a specific class, instance segmentation assigns all voxels belonging to the same object a unique label. Since those labels can not be predicted directly, alternative local representations are sought, which permit extraction of globally unique labels in a subsequent processing step.

The most straight forward local representation is to label pixels as either foreground or background, and then perform a connected component analysis limited to foreground pixels to extract unique objects. However, in the case of 3D neuron segmentation this approach often fails to distinguish voxels in finer neurites, where the axial resolution of the data is lower (Ciresan et al., 2012). To deal with those situations, several methods have centered around the prediction of affinities (*i.e*., the labeling of edges between neighboring voxels as “connected” or “cut”), rather than labeling the voxels themselves (Turaga et al., 2010; Funke et al., 2019).

Affinities effectively increase the resolution of the prediction, but otherwise inherit the advantages and disadvantages of voxel-wise boundary labeling: Both can be computed locally during training and inference, which allows for trivial parallelization. However, segmentations extracted from those predictions are sensitive to small errors: A few incorrectly assigned voxels (or edges between voxels) can label a boundary as foreground, resulting in two segments becoming falsely merged during post-processing.

Those so-called merge errors are the notorious failure modes of neuron segmentation methods. Merge errors have generally been considered worse than split errors, since they have the potential to propagate throughout a dataset. Even if just two neurons are merged together, the resulting segmentation can be difficult to resolve for proofreaders, if the neurons in question have several contact sites. This has been a particular concern for the first generation of proof-reading tools, which did not provide algorithmic help to split wrongly merged objects. In those situations, the origin point of a false merge would first need to be found before the objects can be separated; a task akin to searching for a needle in a haystack.

To avoid those merge errors, Turaga et al. (2009) introduced Malis, a loss function that penalizes topological errors by minimizing the Rand Index. The Rand Index naturally favors split errors over merge errors and thus helped to bias boundary predictions to split instead of merge in ambiguous situations. Funke et al. (2019) expanded this method by constraining the loss to a positive and negative pass, and by providing a maximal spanning tree formulation of the loss, which allows for a quasi-linear and exact computation of the loss during training.

More recent methods do not explicitly focus on merge errors, which is a possible consequence of improved proofreading tools that allow users to separate objects with just a few interactions. Lee et al. (2017) found that using an increased affinity neighborhood acts as an auxiliary learning objective to improve direct neighbor affinities. This work demonstrates that auxiliary learning helps to make better use of local context in the receptive field of the neural network. The nature of this auxiliary learning approach is similar to the LSDs proposed here.

All affinity-based methods have in common that they need a subsequent agglomeration step to produce a final segmentation. Methods such as watershed variants (Wolf et al., 2018) and constrained agglomeration (Beier et al., 2017) successfully demonstrated an increase in robustness of the resulting segmentation to small errors in the predicted affinities.

Not all neuron segmentation methods are based on boundary predictions. The most notable exception are Flood-Filling Networks (FFN), the current state of the art in terms of segmentation quality (Januszewski et al., 2018). FFN eliminated the need for a multi-step segmentation process by using a recurrent convolutional neural network to fill neurons iteratively in an end-to-end fashion. Given seed points within neurons, the algorithm predicts which voxels belong to the same object as the seeds. This approach has been proven to be successful on very large volumes, although it is computationally more expensive than its affinity-based counterparts.

More recently, another promising alternative to boundary prediction has been proposed by Lee et al. (2019), which uses metric learning to produce dense voxel embeddings. The embeddings of voxels that belong to the same object are encouraged to be close in embedding space, while the embeddings of voxels of different objects are pushed away from each other. Clustering of the embeddings then reveals a segmentation. Since object similarity or dissimilarity can only be discerned locally, the method is applied in a block-wise fashion and the segmentations of neighboring blocks are stitched together to process a whole volume.

### 1.2 Contributions

We introduce LSDs as an auxiliary learning task for affinity predictions and demonstrate that segmentation results are competitive with the current state of the art, albeit two orders of magnitude more efficient to compute. LSDs are 10-dimensional vectors, computed for each voxel, which encode local object properties. We engineered LSDs to describe features that could be leveraged to improve boundary detection. Specifically, they consist of three parts: the local size of the object (1D), the offset to the local center of mass (3D), and the local directionality (6D) (described in detail in Section 2.1).

We conducted a large comparative study of recent neuron segmentation algorithms. Specifically, we evaluated four of the aforementioned methods against three LSD-based methods on the following datasets:

- **Z****ebrafinch**: A region consisting of ∼10^6^µm^3^ (∼663 gigavoxels) of songbird neural tissue, imaged using serial block-face EM at 9×9×20 nanometer (xyz) resolution (Januszewski et al., 2018). 0.02% of the full dataset was used to train networks (dense segmentations), 12 manually traced neuron skeletons (13.5 mm) were used for validation and 50 skeletons (97 mm) were used for testing.
- **H****emi-brain**: Three volumes containing ∼1650µm^3^, ∼4750µm^3^, and ∼10360µm^3^ (∼33 total gigavoxels) of raw data, cropped from the ~26 teravoxel dataset generated by Scheffer et al. (2020). This volume was taken from the central brain of a *Drosophila* and imaged with Fib-SEM at 8 nanometer isotropic resolution. 0.002% of the data was used for training (dense segmentation), and 0.06% for testing using a whitelist of proofread neurons.
- **F****ib****-25:** A ∼1.8 10^5^µm^3^ (∼346 gigavoxels) volume from the *Drosophila* visual system was imaged with Fib-SEM at 8 nm isotropic resolution (Takemura et al., 2015). 0.09% of the data was used for training (dense segmentation). Testing was restricted to a 13.8 gigavoxel region using a whitelist of proofread neurons.

For each dataset, we compare LSD-based methods against three previous affinity-based methods: (1) direct neighbor and (2) long-range affinities with mean squared error (MSE) loss, and (3) direct neighbor affinities with Malis loss. Each affinity-based method (including our LSD methods) was trained in the same way and uses the same network architecture (where possible). We used the same segmentation extraction method (from Funke et al. (2019)) to convert the predicted affinities into segmentations.

We further include a comparison against FFN segmentations, which were made available to us by the authors of Januszewski et al. (2018).

We make the training scripts and datasets used in this study publicly available in a central repository^1^, in the hope that non-affinity-based methods that we did not cover in this study (like the recent deep metric learning proposed in Lee et al. (2019)) can be compared in a similar manner.

We compare the aforementioned methods against three different architectures that use LSDs as an auxiliary loss: a simple multitask approach (MtLsd) and two auto-context approaches (AcLsd and AcRLsd). We summarize those methods briefly in the following, for a detailed description see Section 2.2.

MtLsd uses a similar strategy to the long-range affinity neighborhood proposed by Lee et al. (2017). The network is taught to simultaneously predict LSDs and affinities (Figure 1.E).

**Figure 1:**
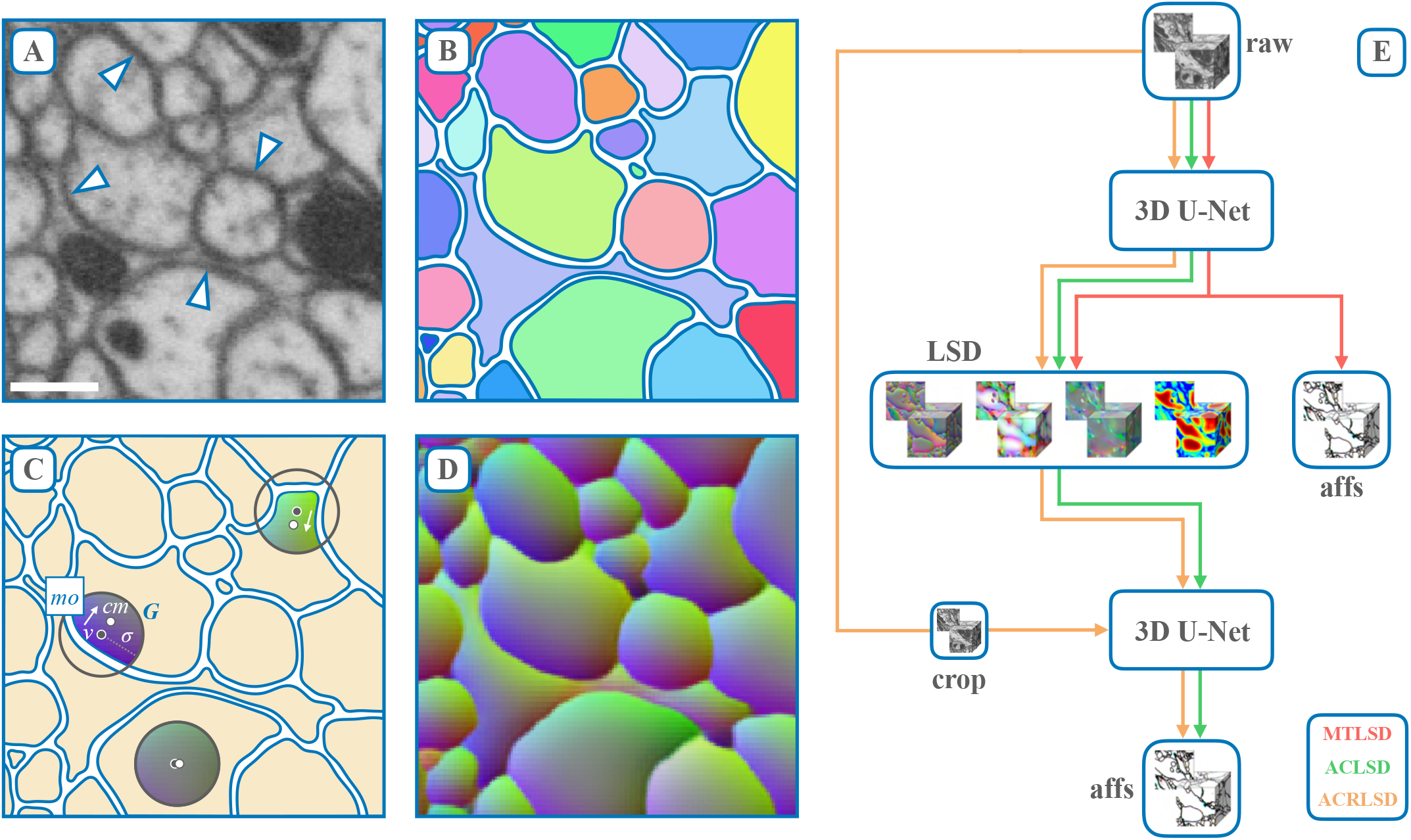
Local shape descriptor and network architecture overview. **A**. EM data imaged with Fib-SEM at 8 nm isotropic resolution (Hemi-brain dataset). Arrows point to example individual neuron plasma membranes. Dark blobs are mitochondria. Scale bar = 300 nm. **B**. Label colors correspond to unique neurons. **C**. LSD mean offset schematic. A Gaussian (G) with fixed sigma (*σ*) is centered at voxel (v). The Gaussian is then intersected with the underlying label (colored region) and the center of mass of the intersection (cm) is computed. The mean offset (mo) between the given voxel and center of mass is calulated (among several other statistics, see Figure 2 for a visualization), resulting in the first three components of the LSD for voxel (v). **D**. Predicted mean offset component of LSDs (Lsd[0:3]) for all voxels. A smooth gradient is maintained within objects while sharp contrasts are observed across boundaries. 3D vectors are RGB color encoded. **E**. Network architectures used. The 10D LSD embedding is used as an auxiliary learning task for improving affinities. In a multi-task approach (MtLsD), LSDs and affinities are directly learnt. In an auto-context approach, the predicted LSDs are used as input to a second network in order to generate affinities both without raw data (AcLsd), and with raw data (AcRLsd). For a detailed network architecture visualization, see Supplementary Figure 11.

AcLsd and AcRLsd use an auto-context learning strategy, as proposed by Tu and Bai (2010). This strategy attempts to refine the quality of a prediction by using a cascade of predictors. For voxel classification, for example, the first pass of an auto-context classifier predicts voxel labels from raw data. The second pass then uses those predictions from the first pass as input^2^. We loosely adapted this idea when designing our auto-context networks. We first taught a network to predict LSDs from raw EM data. The predicted LSDs were then passed into a second network in order to learn affinities (Figure 1.E).

We generally observe an increase in affinity prediction accuracy when training to predict LSDs in an auxiliary task. This increase is most noticeable when using an auto-context setup. We evaluated all methods using two commonly used neuron segmentation accuracy metrics, Variation of Information (VOI) and Expected Run-Length (ERL). We generally find on-par performance between LSD-based methods and FFN under VOI, but inferior scores for LSD-based methods under ERL.

We further investigated how VOI and ERL relate to proof-reading effort given the capabilities of contemporary proof-reading tools (Zhao et al., 2018; Dorkenwald et al., 2022) and found that VOI is a considerably more reliable proxy for proof-reading effort than ERL (see Figure 3B, 3D, Figure 18, and Section 4.1). To this end, we developed a novel metric, the Min-Cut Metric (MCM), to count the number of split and merge operations needed to correct an automatic segmentation. We find general ranking agreement between VOI and MCM, but not between ERL and MCM, which likely stems from ERL’s sensitivity to merge errors. Our results suggest that VOI can serve as a proxy to measure proof-reading effort.

## 2 Methods

### 2.1 Local Shape Descriptors

The motivation behind Local Shape Descriptors (LSDs)^3^ is to provide an auxiliary learning task that improves boundary prediction by learning statistics describing the local shape of the object close to the boundary. A similar technique was already shown to yield superior results over boundary prediction alone (Bai and Urtasun, 2017). Here, we extend on this idea by predicting for every voxel not just affinities values to neighboring voxels, but also statistics extracted from the object under the voxel aggregated over a local window, specifically: (1) the volume, (2) the voxel-relative center of mass, and (3) pairwise coordinate correlations. See Figure 1 for a visualization.

Intuitively, the LSD components encourage the neural network to make use of its entire field of view (FOV) to reach a decision about the presence or absence of a boundary in the center of the FOV. Trained on a boundary prediction task alone (*i.e*., pure affinity-based methods), a neural network might focus only on a few center voxels to detect membranes and achieve high accuracy during training, especially if trained using a voxel-wise loss. However, this strategy might fail in rare cases where boundary evidence is ambiguous. Those rare cases contribute little to the training loss, but given the large size of datasets in connectomics, those cases still result in many topological errors during inference. If, however, the network is also tasked to predict the local statistics of the objects surrounding the membrane, focusing merely on the center voxels is no longer sufficient. Instead, the network will have to make use of its entire field of view to predict those statistics. We hypothesize that this leads to more robust internal representations of objects, allowing the network to infer membrane presence from context, even if the local evidence is weak or missing. Many local object statistics are conceivable that would incentivise the network to use its entire FOV. Here, we focus on simple statistics that are efficient to compute during training.

More formally, let Ω ⊂ ℕ^3^ be the set of voxels in a volume and y : Ω ↦ {0, …, *l*} a ground-truth segmentation. A segmentation induces ground-truth affinity values aff 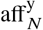, defined on a voxel-centered neighborhood *N* ⊂ ℤ, *i.e*.:

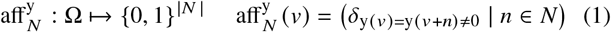

where *δ* is the Kronecker function, *i.e*., *δ* _*p*_ = 1 if predicate *p* is true and 0 otherwise. Our primary learning objective is to infer affinities from raw data x : Ω ↦ ℝ, *i.e*., we are interested in learning a function

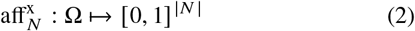

such that 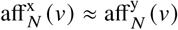.

Similarly to the affinities, we introduce a function to describe the local shape of a segment *i* ∈ {1, …, *l*} under a given voxel *ν*. To this end, we intersect the segment y(*ν*) underlying a voxel *ν* ∈ Ω with a 3D ball of radius *σ* centered at *ν* to obtain a subset of voxels *S*_*ν*_ ⊂ Ω, formally given as

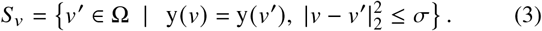

We describe the shape of *S*_*ν*_ by its size, mean coordinates, and the covariance of its coordinates, *i.e*.:

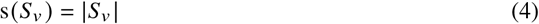

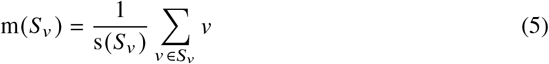

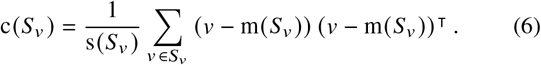

The local shape descriptor lsd^y^ : Ω ↦ ℝ^10^ for a voxel *ν* is a concatenation of the size, center offset, and coordinate covariance, *i.e*.,

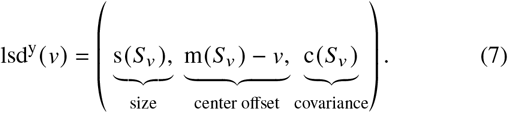

For a visual representation of the components of the LSDs, see Figure 2. We use lsd^y^(*ν*) to formulate an auxiliary learning task that complements the prediction of affinities. For that, we use the same neural network to simultaneously learn the functions aff^x^ : Ω ↦ [0, 1]^|*N*|^ and lsd^x^ : Ω ↦ ℝ^10^ directly from raw data x, sharing all but the last convolutional layer of the network.

**Figure 2:**
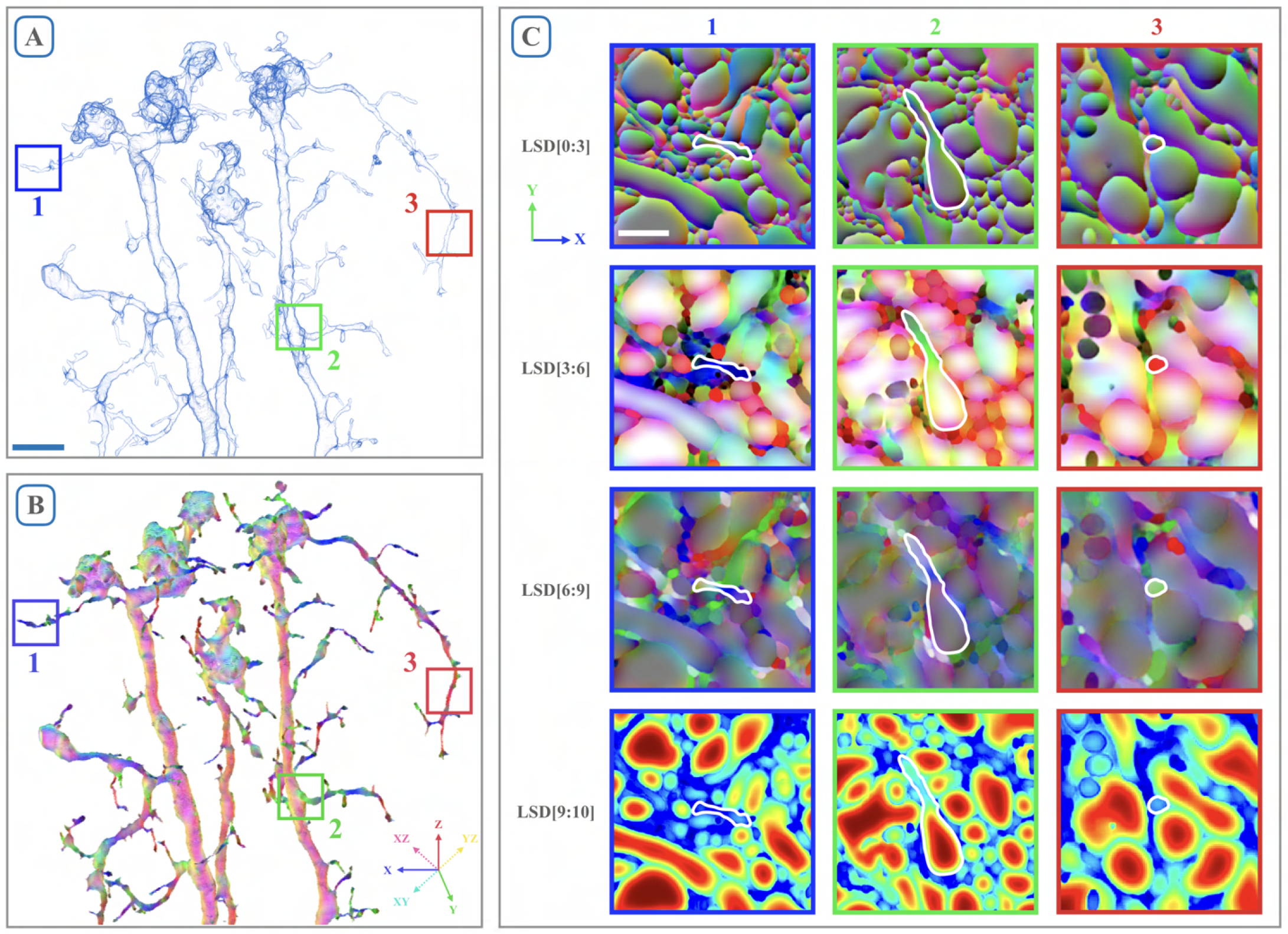
Visualization of LSD components. **A**. Surface mesh of a segmented neuron from Fib-SEM data. Scale bar = 1 µm. **B**. RGB mapping of LSD components 3, 4, and 5. Neural processes are colored with respect to the directions they travel. Intermediate directions are mapped accordingly (see cartesian coordinate inset). **C**. LSD predictions in corresponding 2D slices to the three boxes shown in A and B; neuron highlighted in white. Columns signify neuron orientation (blue = lateral movement, green = vertical movement, red = through-plane movement). Rows correspond to components of the LSDs. First row = mean offset (as seen in Figure 1). Second and third rows = covariance of coordinates (LSD[3:6] for the diagonal entries, LSD[6:9] for the off-diagonals). Second row shows mapping seen in B. Last row = size (number of voxels inside intersected Gaussian). Scale bar = 250 nm.

For efficient computation of the target LSDs during training, the statistics above can be implemented as convolution operations with a kernel representing the 3D ball: Let b^*i*^ : Ω ↦ {0, 1} with b^*i*^ *ν* = *δ*_y(*ν*)=*i*_ be the binary mask for segment *i* and w : ℤ^3^↦ ℝ kernel acting as a local window (*e.g*., a binary representation of a ball centered at the origin, 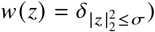. The aggregation of this mask over the window yields the local size s^*i*^(*ν*) of segment *i* at position *ν*. Formally, this operation is equal to a convolution of the binary mask with the local window:

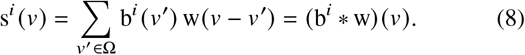

To capture the mean and covariance of coordinates as defined above, we further introduce the following voxel-wise functions m and c. Those functions aggregate the pixel coordinates *ν* over the local window w to compute the local center of mass m^*i*^ (*ν*) and the local covariance of voxel coordinates c^*i*^ (*ν*) for a given segment *i*:

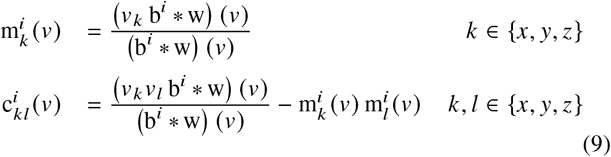

A derivation of those equations can be found in Supplementary Section A.

To obtain a dense volume of shape descriptors, we compute the above statistics for each voxel with respect to the segment this voxel belongs to. Formally, we evaluate for each voxel *ν*:

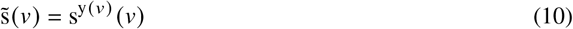

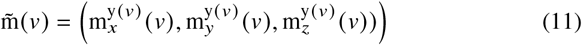

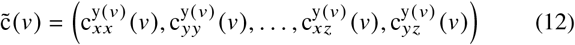

to obtain an equivalent formulation

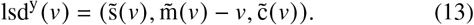

### Network Architectures

We implement the LSDs using three network architectures. The first is a multitask approach, MtLsd, in which the LSDs are output from a 3D U-Net (Çiçek et al., 2016), along with nearest neighbor affinities in a single pass. The other two methods, AcLsd and AcRLsd are both auto-context setups in which the LSDs from one U-Net are fed into a second U-Net in order to produce the affinities. The former relies solely on the LSDs while the latter also sees the raw data in the second pass. For a visualization of network architectures, see Figure 1.E and Supplementary Figure 11. All training was done using Gunpowder^4^ and Tensorflow^5^, using the same 3D U-Net architecture (Funke et al., 2019).

### 2.3 Post-Processing

Prediction and post-processing (*i.e*., watershed and agglomeration) was done in a block-wise fashion using the method described in Funke et al. (2019).

## 3 Results

In this section, we present experimental results of the proposed LSDs for neuron segmentation. We compare the accuracy of the segmentations against several alternative methods for affinity prediction and FFN on three large and diverse datasets. Furthermore, we compare the computational efficiency of different methods and analyze the relationship between different error metrics for neuron segmentations.

### 3.1 Methods

For each dataset we investigated seven methods:

- **Direct neighbor affinities (Baseline):** Baseline network with a single voxel affinity neighborhood and MSE loss as proposed by Turaga et al. (2010). We trained a 3D U-Net to predict affinities.
- **Long range affinities (LR):** Same approach as the Baseline network, but uses an extended affinity neighborhood with three extra neighbors per direction, as proposed by Lee et al. (2017). The extended neighborhood functions as an auxiliary learning task that was shown to improve the direct neighbor affinities.
- **M****alis** **loss (Malis):** Same approach as the Baseline network, but using the loss described in Funke et al. (2019) instead of plain MSE.
- **Flood filling networks (FFN):** A single segmentation per investigated dataset, provided by the authors of Januszewski et al. (2018).
- **Multi-task LSDs (MtLsd):** A network to predict both LSDs and direct neighbor affinities in a single pass, as seen in Figure 1.E. Similar to LR, the LSDs act as an auxiliary learning task for the direct neighbor affinities.
- **Auto-context LSDs (AcLsd):** An auto-context setup, where LSDs were predicted from one network and then used as input to a second network in which affinities were predicted.
- **Auto-context LSDs with raw (AcRLsd):** Same approach as AcLsd, but the second network also receives the raw data as input in addition to the LSDs generated by the first network.

All network architectures are described in detail in Supplementary Section C.1.2 and Supplementary Section D.1.2 for the Zebrafinch and FIB-SEM volumes, respectively.

In order to fairly evaluate accuracy as a function of only the used segmentation method, we made sure to hold other contributing factors constant. Each affinity-based network was trained with the same pipeline (*e.g*. data augmentations and optimizer) and same hyper-parameters for each dataset. We also used the masks to restrict segmentation and evaluation to dense neuropil. These are the same masks used by FFN, thus comparing pure neuron segmentation performance of each method.

### 3.2 Metrics for Neuron Segmentation

Since proofreading of segmentation errors is currently the main bottleneck in obtaining a connectome (Scheffer et al., 2020), metrics to assess neuron segmentation quality should ideally reflect the time needed for proofreading. This requirement is not easily met, since it depends on the tools and strategies used in a proofreading workflow. Currently used metrics aim to correlate scores with the time needed to correct errors based on assumptions about the gravity of certain types of errors. A common assumption has been that false merges take significantly more time to correct than false splits, although next generation proofreading tools challenge this conception (Plaza and Funke, 2018).

In this study, we report neuron segmentation quality with two established metrics: Variation of Information (VoI) and Expected Run-Length (ERL). In addition to those metrics, we propose a new metric, which we refer to as the Min-Cut Metric (MCM), designed to measure the number of graph edit operations needed to perform in a hypothetical proofreading tool.

#### 3.2.1 Variation of Information (VoI)

A metric to compare clusterings (Meilă, 2007), which became an established metric to assess neuron segmentation accuracy. VoI measures the disagreement between two segmentations in terms of the average number of bits needed to guess the segment ID of a randomly chosen voxel in one segmentation, given only its label in the other segmentation. This measurement is performed in both directions, giving rise to the two additive components of VoI, a measure for split and merge errors. Lower values are better, with equivalent segmentations (up to label permutations) having a value of zero.

#### 3.2.2 Expected Run-Length (ERL)

Following the assumption that false merges are in practice harder to correct than false splits, Januszewski et al. (2018) proposed to measure accuracy in terms of the Expected Run Length (ERL), which measures the expected length of an error-free path along neurons in a volume. Notably, all paths contained in falsely merged segments are considered erroneous, thus ERL emphasizes merge errors disproportionally. An appealing aspect of ERL is that it relates segmentation errors to cable length, a commonly used and interpretable feature of neurons.

#### 3.2.3 Min-Cut Metric (MCM)

A metric that assumes that a user can directly interact with agglomerated fragments from an oversegmentation. In particular, we assume that users can split segments by means of a mincut through the fragment graph between two selected fragments, where edge costs correspond to the merge scores used during agglomeration. In this context, the MCM reports the number of split and merge operations needed to be performed by a human annotator to obtain the desired segmentation.

Notably, a merge of two neurons can be resolved with a single interaction, even if the neurons have several contact sites. The min-cut solution will identify all necessary cuts to separate the two selected fragments. For most commonly used agglomeration algorithms (and the one used here), only one cut is necessary, as connectivity on the fragment graph is defined by a single linkage clustering (Funke et al., 2019).

This metric is of practical relevance since min-cuts on fragment graphs are used to split merged objects in commonly used proofreading tools (Zhao et al., 2018; Dorkenwald et al., 2022). The details of this metric are described in Supplementary Section B.

### 3.3 Datasets

The following describes the datasets, regions of interest (RoIs), and ground-truth used in this study. Dataset overviews can be seen in Figure 3, Figure 4, Figure 5 top panels, and are described in Table 1.

**Figure 3:**
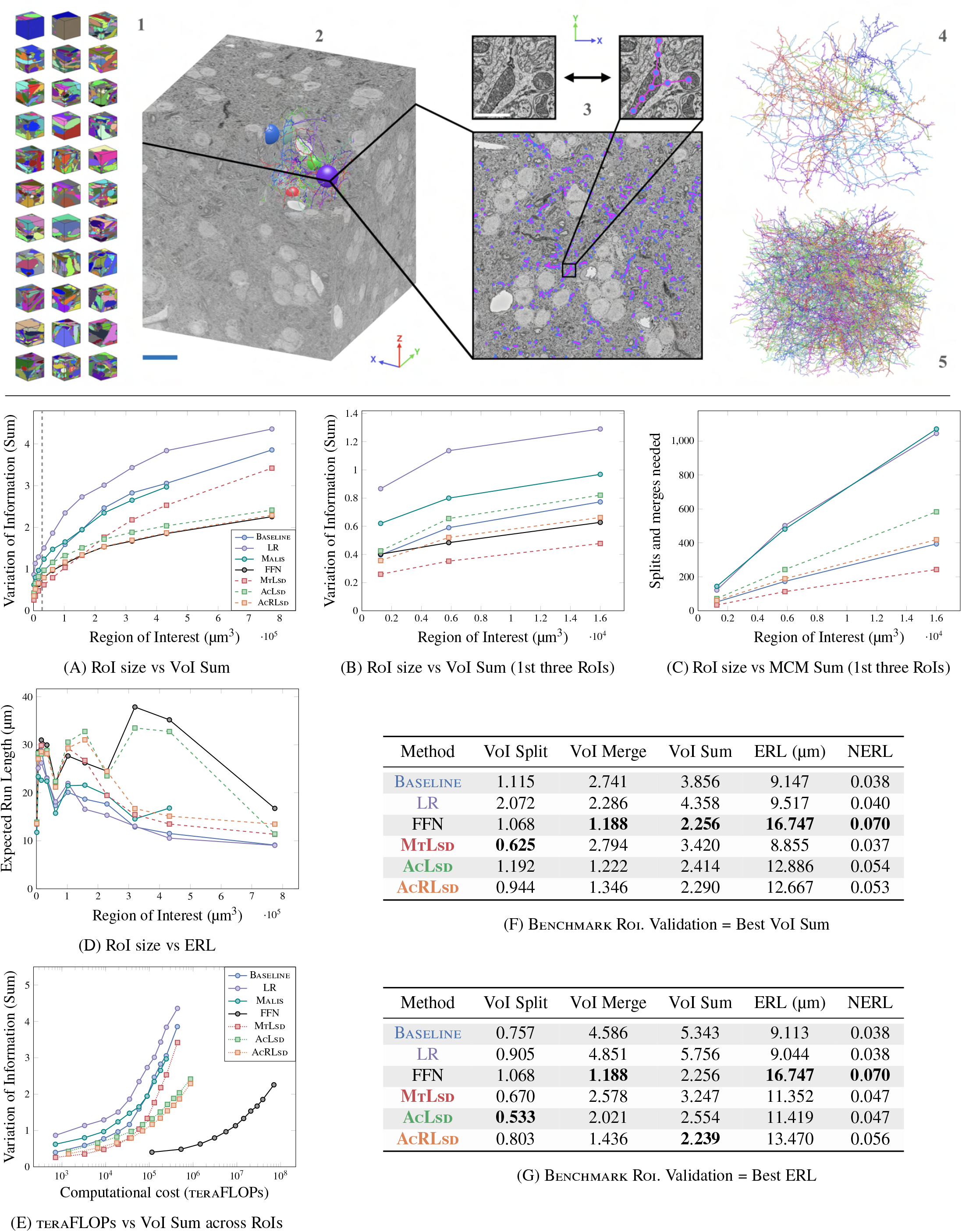
Zebrafinch dataset. Top panel shows data used for training and testing. **1**. 33 gound truth volumes were used for training. **2**. Full raw dataset, scale bar = 15 µm. **3**. Single section shows ground-truth skeletons, zoom in scale bar = 500 nm. **4**. Validation skeletons (n=12). **5**. Testing skeletons (n=50). Main results shown below top panel. Points in Zebrafinch plots correspond to optimal thresholds from validation set. Each point represents an RoI. For VoI and MCM, lower scores are better, for ERL higher scores are better. **A**. VoI Sum vs RoI size (µm^3^). **B**,**C**. VoI Sum and MCM Sum vs RoI size (first three RoIs), respectively. Dashed line in **A** corresponds to RoIs shown in **B**,**C. D**. ERL (nanometers) vs RoI size. **E**. teraFLOPS vs VoI Sum across RoIs (same as in panels A and D). Tables (**F**,**G**) show best network scores in bold, with respect to which metric was optimized during validation.

**Table 1:**
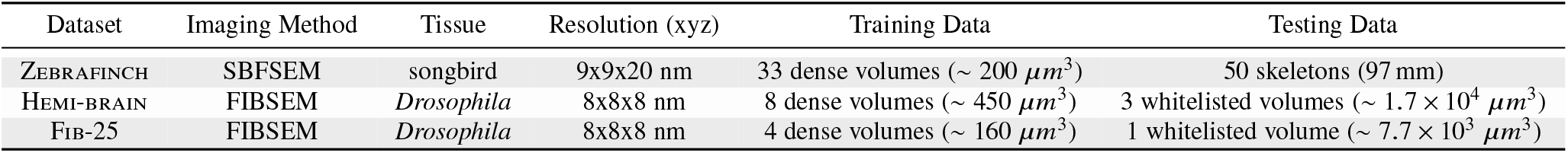
Overview of datasets used in study

**Figure 4:**
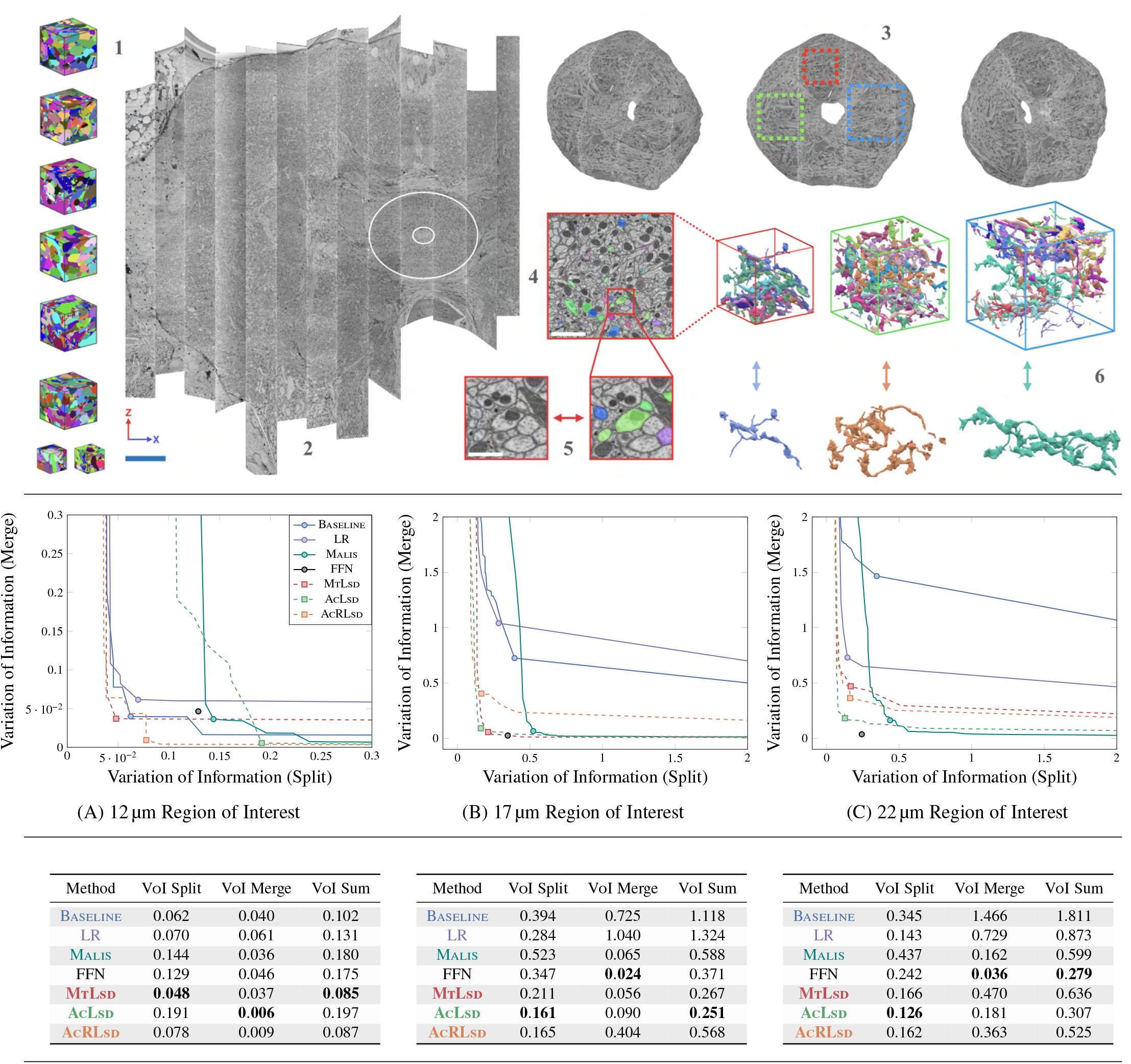
Hemi-brain dataset. Top panel shows data used for training and testing. **1**. 8 ground-truth volumes were used for training. **2**. Full Hemi-brain volume, scale bar = 30 µm. Experiments were restricted to Ellipsoid Body (circled region). **3**. Volumes used for testing. **4**. Example sparse ground-truth testing data, scale bar = 2.5 µm. **5**. Zoom-in, scale bar = 800 nm. **6**. Example 3D renderings of selected neurons. Main results below top panel. Plot curves show results over range of thresholds for each RoI (**A** = 12 µm RoI, **B** = 17 µm RoI, **C** = 22 µm RoI). Points correspond to optimal thresholds on testing set, no validation set was available. Lower scores are better. Tables show best network scores in bold for each RoI.

**Figure 5:**
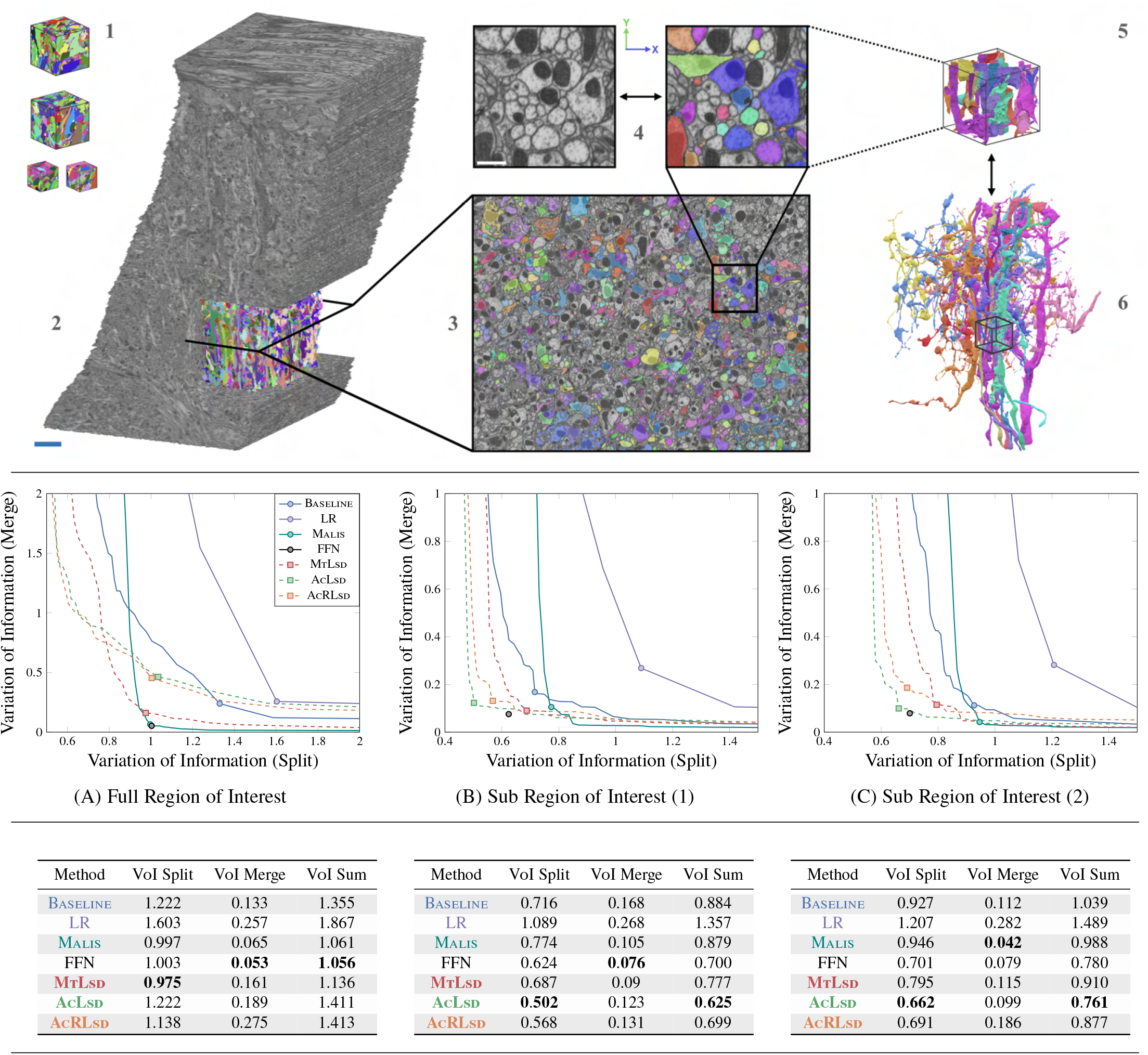
Fib-25 dataset. Top panel shows data used for training and testing. **1**. 4 ground-truth volumes were used for training. **2**. Full volume with cutout showing testing region, scale bar = 5 µm. **3**. Cross section with sparsely labeled testing ground-truth. **4**. Zoom-in, scale bar = 750 nm. **5**. Sub-volume corresponding to zoomed-in plane. **6**. Subset of full RoI testing neurons. Small volume shown for context. Main results below top panel. **A**. Full testing RoI. **B**,**C**. Two sub RoIs contained within full RoI. Plot curves show results over range of thresholds. Points correspond to optimal thresholds on testing set, no validation set was available. Lower scores are better. Tables show best network scores in bold, for each RoI.

**Figure 6:**
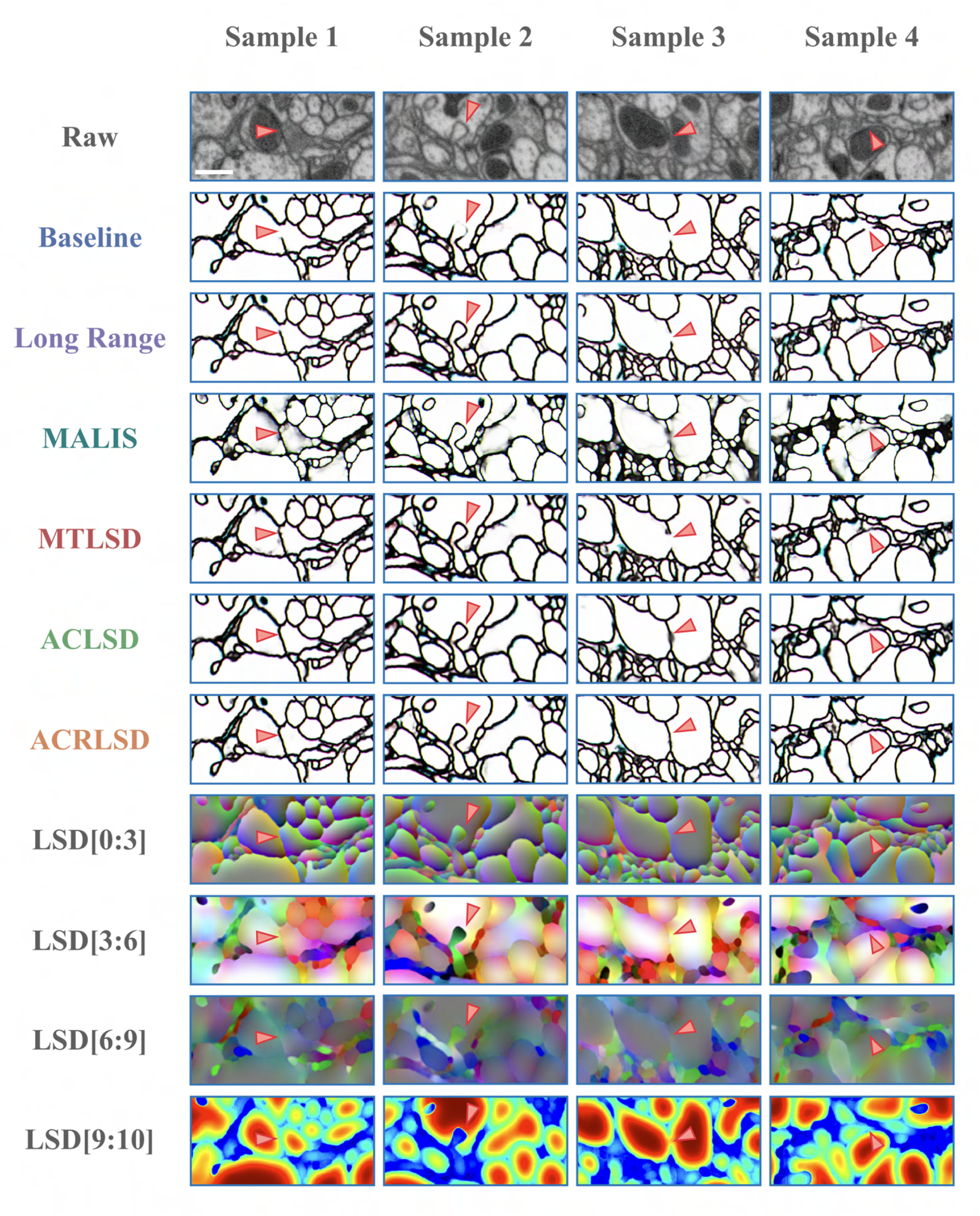
Qualitative examples showing predicted affinities and LSD components on Fib-25 dataset. Scale bar = 500 nm. Top row shows raw data. Arrows correspond to example plasma membranes. A baseline architecture (row two) fails to distinguish these boundaries, which could lead to possible errors in a resulting segmentation. The LSDs (bottom 4 rows) help to improve the affinites generated by the MtLsD, AcLsd and AcRLsd networks.

#### 3.3.1 Zebrafinch

The largest dataset used in this study was the songbird dataset also used in Januszewski et al. (2018). This volume consists of a ∼10^6^µm^3^ region of a zebrafinch brain, imaged with serial blockface EM at a resolution of 9×9×20 nm (x,y,z). For our experiments, we used a slightly smaller region completely contained inside the raw data with edge lengths of 87.3, 83.7, and 106 µm (x,y,z). We refer to this region as the Benchmark Roi.

For each affinity-based network described in Section 3.1, we used 33 volumes containing a total of ∼200µm^3^ (∼6µm^3^ average per volume) of labeled data6 for training. We then ran prediction on the Benchmark Roi, using a block-wise processing scheme.

Using the resulting affinities, we generated two sets of supervoxels: one without any masking and one constrained to neuropil using a mask^6^. Additionally, we filtered supervoxels in regions in which the average affinites were lower than a predefined value (*e.g*., glia). Supervoxels were agglomerated using one of two merge functions, described in Funke et al. (2019), to produce the region adjacency graphs used for evaluation.

We then produced segmentations for RoIs of varying size centered in the Benchmark Roi, in order to assess how segmentation measures scale with volume size. In total, we cropped 10 cubic RoIs ranging from ∼11µm to ∼76µm edge lengths, in addition to the whole Benchmark Roi. We will refer to the respective RoIs by their edge lengths. For each affinity-based network, in each RoI, we created segmentations for a range of agglomeration thresholds (resulting in a sequence of segmentations ranging from over- to undersegmentation). Additionally, we cropped the provided FFN segmentation accordingly and relabeled connected components.

We used a set of 50 manually ground-truthed skeletons^6^, comprising 97 mm, for evaluation. For each network we assessed VoI and ERL on each RoI. For affinity-based methods we also computed the MCM on the 11, 18, and 25µm RoIs. Additionally, we used 12 validation skeletons consisting of 13.5 mm to determine the optimal thresholds for each network on the Benchmark Roi. Further details can be found in Supplementary Section C.

#### 3.3.2 Hemi-Brain

The so-called Hemi-brain is a Fib-SEM volume of the *Drosophila melanogaster* central brain, imaged at 8nm isotropic resolution (Scheffer et al., 2020), comprising a total of 26 teravoxels of image data. We evaluate all investigated methods on regions restricted to the Ellipsoid Body, a neuropil implicated in spatial navigation, (Turner-Evans and Jayaraman, 2016), which contained ample ground-truth data for evaluation.

We used eight volumes of densely annotated ground-truth volumes containing ∼ 450µm^3^ of labeled data for training. Three RoIs with ∼ 12µm, ∼ 17µm, and ∼ 22µm edge lengths were cropped from the larger volume, and prediction was done directly on each RoI.

Supervoxels were limited to the Ellipsoid Body using a mask7 and then agglomerated using the same two merge functions as in the Zebrafinch dataset.

We produced segmentations for each network over a range of thresholds on the RoIs, and consolidated a single FFN segmentation^6^. A densely labeled ground-truth volume was cropped and filtered using a list of neuron IDs^7^ deemed to be completely traced by expert proofreaders. Since the ground-truth is comprised of voxel data rather than skeletons, we report only VoI. Further details can be found in Supplementary Section D.

#### 3.3.3 FIB-25

The Fib-25 dataset, produced by Takemura et al. (2015), is another Fib-SEM volume imaged at 8nm resolution, containing ∼1.8 × 10^5^µm^3^ of raw data taken from the *Drosophila* visual system. Four volumes with ∼ 160µm^3^ of labeled data^7^ were used for training.

We predicted on the full raw data and then restricted supervoxels to an irregularly shaped neuropil mask^6^. Agglomeration was done in the same fashion as the aforementioned volumes. We created segmentations in two ways: for the first method, following the procedure described in Januszewski et al. (2018), we segmented neurons within the entire neuropil mask. For the second method, we limited segmentation to two sub-RoIs contained within both the neuropil mask and testing region (sections 5074–7950). The sub-RoIs have a size of ∼2.2 × 10^3^µm^3^ and ∼ × 2.6 10^3^µm^3^, respectively. For FFN, we cropped the provided segmentation^6^ and relabeled connected components.

Evaluation was limited to a list of proofread, voxel ground-truth labels^7^ contained inside the testing region. Depending on the segmentation method, connected components on both ground-truth and segmentation volumes were either cropped and relabelled or untouched. Since the ground-truth is comprised of voxel data rather than skeletons, we report only VoI. Further details can be found in Supplementary Section D.

### 3.4 Neuron Segmentation Accuracy

#### 3.4.1 Zebrafinch

We find that LSDs are useful for improving the accuracy of direct neighbor affinities, and subsequently the resulting segmentations. Specifically, LSD-based methods consistently outperform other affinity-based methods over a range of RoIs, whether used in a multitask (MtLsd) or auto-context (AcLsd and AcRLsd) architecture (Figure 3.A). In terms of segmentation accuracy according to VoI, the best auto-context network (AcRLsd) performs on par with FFN (Figure 3.A).

Interestingly, we find that the ranking of methods depends on the size of the evaluation RoI. Even for monotonic metrics like VoI, we see that performance on the smallest RoIs (up to 54 µm) does not extrapolate to the performance on larger datasets.

We also investigated how ERL varies over different RoI sizes. To this end, we cropped the skeleton ground-truth to the respective RoIs and relabeled connected components (as we did for the VoI evaluation). However, the resulting fragmentation of skeletons heavily impacts ERL scores: ERL can not exceed the average length of skeletons, and thus the addition of shorter skeletons fragments can result in a decrease of ERL, even in the absence of errors. We see this effect prominently in Figure 3.D: ERL measures do not progress monotonically over RoI sizes and absolute values are likely not comparable across different dataset sizes. In addition, the ranking of methods for a given RoI size varies significantly over different RoI sizes. The discrepancy between the computed ERL and maximum possible ERL (or the ground truth ERL) further emphasizes this point (Figure 10).

Furthermore, the ERL metric is by design very sensitive to merge errors, as it considers a whole neuron to be segmented incorrectly if it was merged with even only a small fragment from another neuron. Thus, merge errors contribute disproportionally to the ERL computation. In addition, the contribution depends on the sizes of the merged segments. Merging a small fragment of one neuron into an otherwise correctly reconstructed large neuron will have a larger negative impact on the ERL than merging two small fragments from different neurons, although the effort needed to resolve that error is likely the same. We observe that this property leads to erratic scores across different volume sizes (see Figure 3.D, Figure 9) that no longer reflect the amount of time needed to proof-read the resulting segmentation. The sensitivity to merge errors also contribute to the observed differences between the ERL scores of the LSD-based methods and FFN (Figure 3D): Although AcRLsd has a lower total VoI of 2.239 than FFN with 2.256, AcRLsdhas a higher merge rate with a VoI merge score of 1.436 than FFN with a VoI merge score of 1.118, resulting in substantially different ERL scores of 13.5µm for AcRLsd and 16.7µm for FFN, (Figure 3G).

The high variability between metrics and RoI sizes prompted us to develop a metric that aims to measure proofreading effort. We developed MCM to count the number of interactions needed to split and merge neurons in order to correctly segment the ground-truth skeletons, assuming that a min-cut-based split tool is available. Due to the computational cost associated with MCM (stemming from repeated min-cuts in large fragment graphs), we limited its computation to the three smallest investigated RoIs in this dataset. As expected, we observe a linear increase in MCM with RoI size across different methods (Figure 3.C). Furthermore, we see that MCM and VoI mostly agree on the ranking of methods (Figure 3.B, Supplementary Figure 8), which suggests that VoI should be preferred to compare segmentation quality in the context of a proofreading workflow that allows annotators to split false merges using a min-cut on the fragment graph. Since the MCM requires a supervoxel graph, it was not possible to compute on the single FFN segmentation provided.

#### 3.4.2 Hemi-Brain

We observe a similar variability of method rankings over RoI sizes on the Hemi-brain dataset.

On the largest investigated RoI (22µm edge length), AcLsd clearly performs best among all affinity-based methods (VoI sum of 0.307 vs. 0.525 for the second best, see Figure 4.C). As such, AcLsd is competitive with FFN (VoI sum 0.279), with the notable difference of performing more merge errors, but significantly less split errors than FFN.

On the 17µm RoI, AcLsd again performs better than all other affinity-based methods and also better than FFN (VoI sum of 0.251 vs. 0.371, see Figure 4.B). MtLsd is on par with AcLsd on this RoI, which stands in contrast to its significantly worse performance on the 22µm RoI. This observation further puts into question to what extent the accuracy of a segmentation can be extrapolated from smaller to larger RoI sizes.

These concerns become even more evident when turning to the results on the smallest RoI of 12µm edge length. Here, AcRLsd performs worse than all other methods, with a significant margin to the best performing method, MTLSD (VoI sum 0.197 vs 0.085). Even Baseline achieves very good results on this RoI (VoI sum 0.102), although it would be a poor choice in production given its detrimental performance on the larger RoIs. We have to conclude that the size of this RoI is likely not large enough to accurately deduce whether the differences in method performance are due to model accuracies or data biases.

A somewhat surprising result is the performance of AcRLsd on this dataset. Although architecturally very similar to AcRLsd (the only difference is that AcRLsd receives raw data and LSDs in the second pass, while AcLsd receives only LSDs), AcRLsd is significantly worse on the two larger RoIs. This stands in contrast to the results we obtained on the Zebrafinch dataset. Our results do not allow us to say with confidence whether this artifact is due to overfitting to the training data (which might be more likely to happen for AcRLsd) or due to model noise introduced by the random initialization during training.

#### 3.4.3 Fib-25

We first evaluated all methods on the full testing RoI of the Fib-25 dataset (Figure 5.A). On the full RoI, LSDs generally do not perform well. We observe that the best auto-context method (AcLsd) performs worse than the Baseline (VoI sum 1.413 vs 1.355). Interestingly, MtLsd achieves higher accuracy than both auto-context networks. FFN exceeds all other methods (VoI sum 1.056)^8^, and Malis is not far behind (VoI sum 1.061). Since those results are not consistent with the results seen on the Zebrafinch and Hemi-brain volumes, we visually inspected the segmentations. We found a high rate of false merges occurring in the periphery of the testing RoI, stemming from nuclei and boundaries of the imaged volume, which are not contained in the training data. As such, the full testing RoI of this dataset favors “conservative” methods, *i.e*., methods that have higher split rates.

To test the plain neuropil segmentation accuracy, we further cropped two RoIs (∼ 2.2 10^3^µm^3^ and ∼ 2.6 10^3^µm^3^) inside the testing region, such that they contain only dense neuropil (see Supplementary Section D.3.3 for details).

On the two sub RoIs, LSDs outperform other affinity-based methods and are comparable to FFN, (Figure 5.B,C). Consistent with the Zebrafinch and Hemi-brain results, using an auto-context approach seems to generate the best results. On sub RoI 1, AcLsd slightly exceeds the accuracy of FFN (VoI sum: 0.625 vs 0.700, respectively). We observe similar results on sub RoI 2 (VoI sum: 0.761 vs 0.780, respectively).

These results highlight the need for masking neuropil when processing large volumes, as was done in the Zebrafinch dataset. Interestingly, LR affinities perform poorly across RoIs, which might suggest that an increased affinity neighborhood is sometimes not sufficient for improving direct neighbor affinities.

### 3.5 Throughput

In addition to being accurate, it is important for neuron segmentation methods to be fast and computationally inexpensive. As described in the introduction, the acquisition size of datasets is growing rapidly and approaches should therefore aim to complement this trajectory. Since LSDs only add a few extra feature maps to the output of the U-Net, there is almost no difference in computational efficiency compared to Baseline affinities. LSD-based methods can therefore be parallelized in the same manner as affinities, making them a good candidate for the processing of very large datasets or environments with limited computing resources.

In our experiments, prediction and segmentation of affinity-based methods was done in a block-wise fashion, allowing parallel processing across many workers, see Figure 7 for an overview. This allowed for efficient segmentation following prediction (Table 2.B).

**Table 2:**
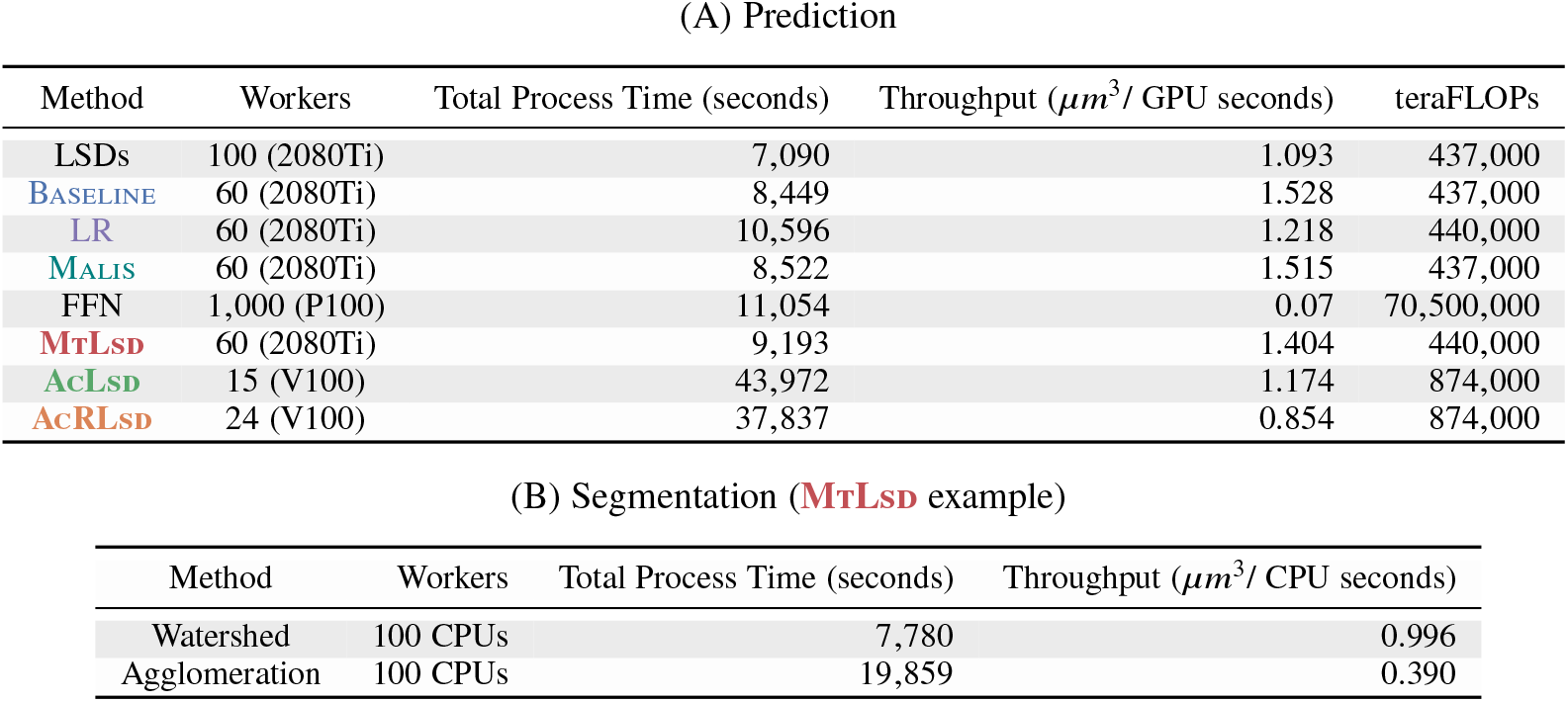
Computational costs on Zebrafinch Benchmark Roi.

**Figure 7:**
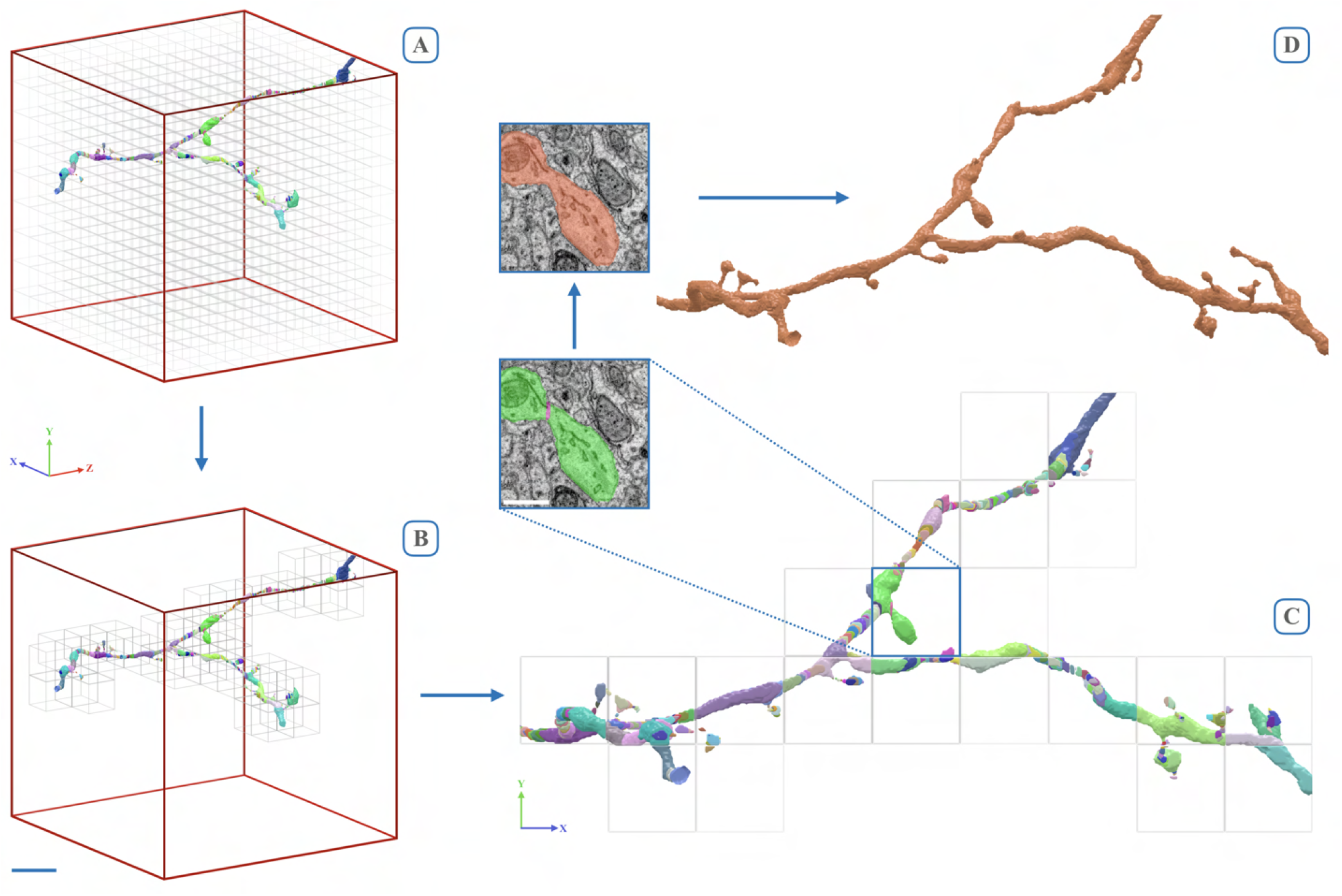
Overview of block-wise processing scheme. **A**. Example 32 µm RoI showing total block grid. **B**. Required blocks to process example neuron. Scale bar = ∼ 6µm. **C**. Corresponding orthographic view highlights supervoxels generated during watershed. Block size = 3.6 µm. Inset shows respective raw data inside single block (scale bar = ∼ 1µm). **D**. Supervoxels are then agglomerated to obtain a resulting segment. **Note:** While this example shows processing of a single neuron, in reality all neurons are processed simultaneosuly.

When considering computational costs in terms of floating point operations (FLOPS), we find that the AcRLsd network (the computationally most expensive of all LSD architectures) is two orders of magnitude more efficient than FFN, while producing a segmentation of comparable quality (Figure 3.E). For this comparison, we computed FLOPS of all affinity-based methods during prediction (see Supplementary Section E for details). For FFN, we used the numbers reported in Januszewski et al. (2018), limited to the forward and backward passes of the network, *i.e*., the equivalent of the prediction pass for affinity-based methods. We limit the computational cost analysis to GPU operations, since FLOP estimates on CPUs are unreliable and the overall throughput is dominated by GPU operations. We therefore only consider inference costs for all affinity-based networks, since agglomeration is a post-processing step done on the CPU. To keep the comparison to FFN fair, we do not count FLOPS during FFN agglomeration, although it involves a significant amount of GPU operations. Table 2.A summarizes the computational costs of all investigated methods and provides throughput numbers on the hardware used in this study. Generally, affinity-based methods are more computationally efficient than FFN by two orders of magnitude.

## 4 Discussion

The main contribution of this work is the introduction of LSDs as an auxiliary learning task for neuron segmentation. We demonstrated that, when compared to other affinity-based methods, the LSDs consistently help to improve neuron segmentations across specimen, resolution, and imaging techniques. We also found results to be competitive with the current state of the art approach, while being two orders of magnitude faster. All methods, datasets, and results are publicly available^9^, which we hope will be a useful starting point for further extensions and a benchmark to evaluate future approaches in a comparable manner.

On the Zebrafinch dataset, the largest dataset both in terms of image data and available ground-truth, LR and Malis did not exceed the accuracy of the Baseline network. This is surprising considering that LR also uses an auxiliary task to improve direct neighbor affinity predictions. This observation might well be due to differences in how masks are handled during training, which we will discuss in more detail in Section 4.4. We found those results generally confirmed on the Hemi-brain dataset, with the exception of Malis, which performed better on this dataset than on Zebrafinch.

Although relatively small in comparison, the Fib-25 dataset nevertheless provided insights into network performance in the periphery of dense neuropil. On the full RoI, both auto-context LSD networks performed very poorly, while Malis was on par with FFN. This is due to increased false merges in the periphery, which proliferate into the testing RoI. Networks that favor splitting over merging, such as Malis and FFN, are consequently less affected. Further analysis on two sub RoIs, both contained entirely within the neuropil mask and the testing RoI, confirmed this to be the case: here, LSD networks improve over other affinity networks and are again competitive with FFN. These results highlight the importance of masking when processing large volumes.

### 4.1 Metric Evaluation

An important but challenging task is finding a robust metric for assessing the quality of a neuron segmentation. Ideally, such a metric reflects the amount of time needed to proofread a segmentation. Here, we presented results in terms of VoI and ERL, two commonly used metrics for this task.

VoI directly reports the amount of split and merge errors. Being a voxel-wise metric, however, VoI can be sensitive to slight, but systematic, shifts in boundaries. At the same time, small topological changes might go unnoticed, which is especially problematic in fine neurites in the vicinity of synapses (Funke et al., 2017).

ERL reports the expected error-free path-length of a reconstruction with respect to skeleton ground-truth. Similar to VoI, ERL is not sensitive to small topological changes close to terminals. Furthermore, ERL disproportionately punishes merge errors and subsequently favors split-preferring methods, explained in Section 3.4.1 (Plaza and Funke, 2018). Additionally, we found that ERL increases non-monotonically with varying volume sizes (Figure 3.D), which is due to the fragmentation of skeletons in volumes that are not large enough to contain entire neurons.

Consequently, neither method directly reflects the labor required for proofreading a segmentation, which is arguably the relevant quantity to optimize (Plaza, 2016; Funke et al., 2017). This quantity depends on the available tools for proofreading, and in particular on the amount of interactions needed to fix errors of different kinds: False splits might be hard to find, but do require only one interaction to merge. False merges, on the other hand, might be easy to spot, but the number of interactions needed to fix them depends greatly on the proofreading tool. Current proofreading tools (Zhao et al., 2018; Dorkenwald et al., 2022) allow annotators to correct merge errors with a few interactions. We therefore introduced the MCM, a metric which uses graph cuts to emulate the amount of interactions required to correct false merges in a segmentation. We observed a linear growth of MCM with volume size (Figure 3.C), which is a necessary condition for neuron segmentation metrics that measure the amount of proofreading effort needed (assuming an equal distribution of errors).

Unfortunately, MCM is computationally quite expensive. The sequence of graph-cuts needed for the evaluation of merge errors quickly becomes infeasible on large volumes. However, MCM shows general ranking agreement with VoI, evaluated on 11 randomly sampled sub-RoIs in Zebrafinch (Supplementary Figure 5) and across different thresholds (Supplementary Figure 8). These findings suggest that VoI can serve as a reasonable proxy to rank methods based on their expected proofreading time.

Additionally, we find VoI to be a robust metric for the validation of method parameters: For each affinity-based network, the threshold minimizing VoI sum on the validation set is also close to the best threshold on the testing set (Supplementary Figure 6). This property is of practical relevance, as in any real-world scenario hyperparameters have to be adjusted on a volume that is significantly smaller than the target volume. Unfortunately, ERL does not seem to exhibit this property to the same degree: the best validation thresholds gradually diverge from the best testing thresholds as scale increases (Supplementary Figure 7), which makes it difficult to extrapolate segmentation accuracy from a validation dataset.

### 4.2 Auxiliary Learning for Boundary Prediction

Auxiliary learning tasks have been shown to improve network performance across different applications. One possible explanation for why auxiliary learning is also helpful for the prediction of neuron boundaries is that the additional task incentivizes the network to consider higher-level features. Predicting LSDs is likely harder than boundaries, since additional local structure of the object has to be considered. Merely detecting an oriented, dark sheet (*e.g*., plasma membranes) is not sufficient; statistics of the whole neural process have to be taken into account. Those statistics rely on features that are not restricted to the boundary in question. Therefore, the network is forced to make use of more information in its receptive field than is necessary for boundary prediction alone. This, in turn, increases robustness to local ambiguities and noise for the prediction of LSDs. As a welcome side-effect, it seems that the network learns to correlate boundary prediction with LSD prediction, which explains why the boundary prediction benefits from using the LSDs as an auxiliary objective.

Surprisingly, we see that LR affinities do not perform as well across the investigated datasets. While LR affinities share some of the benefits of LSDs, they might not be as efficient as LSDs in encoding higher-level features. For example, LR affinities have blind spots (missing neighborhood steps), whereas LSDs are spatially homogeneous. Additionally, we found LR affinities to be detrimental when used with masking of glia and other structures. It is likely harder to correlate nearest neighbor affinities with a long-range neighborhood in the presence of masks.

While Lee et al. (2017) saw superhuman accuracy using an increased affinity neighborhood on the SNEMI3D challenge, the processed volume was relatively small (∼ 110µm^3^). Our results suggest that it is hard to correlate accuracy on small volumes to accuracy on large volumes (Figure 3.A)^10^. Additionally, we only consider an increased affinity neighborhood and not other aspects of the original LR implementation, such as residual modules in the U-Net and inference blending, which might be essential for further performance increases. Finally, the SNEMI3D dataset has an anisotropy factor of ∼ 5, whereas the data we test on here has an anisotropy factor of either ∼ 2 (Zebrafinch) or is isotropic (Hemi-brain, Fib-25).

### 4.3 Auto-Context Refinement

We found that using an auto-context approach greatly improved resulting segmentations (Figure 3.A). We tested if this increase in accuracy was consistent when using affinities as the input to the second network (*i.e*., a Baseline auto-context approach, AcBaseline), and found that it made no significant improvements to the Baseline network (Figure 8).

**Figure 8:**
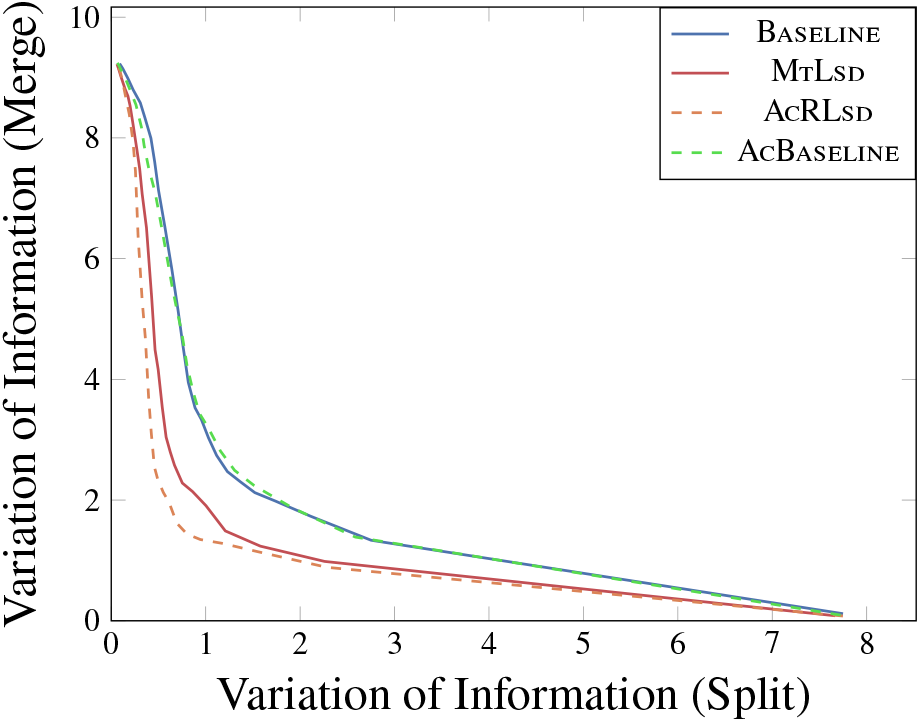
Zebrafinch, Benchmark Roi, VoI split vs VoI merge, auto-context comparison. Using Baseline affinities in an auto-context architecture does not improve accuracy (blue solid and green dashed lines), while the LSDs do benefit (red solid and orange dashed lines).

We hypothesize that predicting affinities from affinities is too similar to predicting affinities from raw EM data. Specifically, we suspect that the AcBaseline network simply copies data in the second pass rather than learning anything new. Easy solutions, such as looking for features like oriented bars, already produce relatively accurate boundaries in the first pass. Consequently, there is little incentive for the network to change course in the second pass. Translating from LSDs to affinities, on the other hand, is a comparatively different task, which forces the network to incorporate the features from the LSDs in the second pass. The subsequent boundary predictions seem to benefit from this.

However, it remains unclear whether auto-context is always necessary. While it does consistently yield optimal results on large datasets, it is likely overkill for small to intermediate sized volumes, due to the increase in computation. Since the MtLsd network still offers improved accuracy over Baseline affinities on every investigated dataset, it is probably sufficient in the majority of cases. In this context, a multi-task network can be considered the default configuration while an auto-context approach should be considered in complex cases where the former would otherwise fail. As always, the optimal solution would ideally be chosen based on an available validation set, as was done for the Zebrafinch.

### 4.4 Masking

Binary masks are commonly used to limit neuron segmentation to dense neuropil and exclude confounding structures like glia cells. Recent approaches to processing large volumes have incorporated tissue masking at various points in the pipeline (Januszewski et al., 2018; Li et al., 2019; Dorkenwald et al., 2019; Scheffer et al., 2020), to prevent errors in areas that were underrepresented in the training data.

Our results confirm the importance of masking. We used a neuropil mask which excluded cell bodies, blood vessels, myelin, and out-of-sample (background) voxels (see Supplementary Figure 15 for a visualization). Across all investigated methods, the accuracy degraded substantially on larger RoIs when processed without masking (see Figure 9, Supplementary Figure 4).

**Figure 9:**
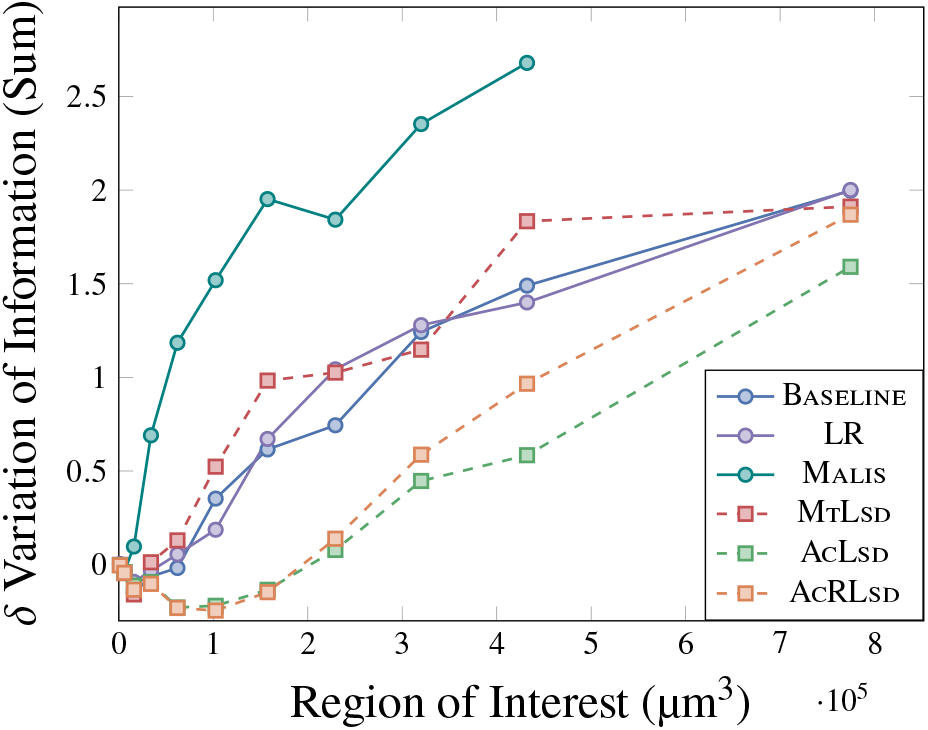
Zebrafinch, mask *δ* VoI sum vs RoI. Accuracy decreases substantially without proper masking as scale increases.

Masking of irrelevant structures can also be incorporated in the training process. The Zebrafinch training volumes already had some glial processes masked out. We trained all networks to predict zero affinities in these regions (see Supplementary Figure 13 for a visualization). We then discarded fragments with close to zero affinity values during agglomeration. Methods which succeeded in learning to mask these areas (Baseline, MtLsd, AcLsd, AcRLsd) produced better results than those that did not (Malis, LR).

### 4.5 Accuracy Extrapolation

One of the challenges of deep learning is to find representative testing data and metrics to infer production performance. This is especially challenging for neuron segmentation, considering the diversity of neural ultrastructure and morphology found in EM volumes. While challenges like CREMI^11^ and SNEMI3D^12^ make an effort to include representative training and testing data, the implications for model performance on larger datasets are not straight forward.

Our results suggest that testing on small volumes provides limited insight into the quality of a method when applied to larger volumes. For example, the total volume of the three CREMI testing datasets (∼1056 µm^3^) is still less than the smallest Zebrafinch (∼ 1260 µm^3^) and Hemi-brain (∼ 1643 µm^3^) RoIs. As shown in Figure 3 and Figure 4, the smallest volumes are not indicative of performance on larger volumes. In this context, it seems difficult to declare a clear “winner” when it comes to neuron segmentation accuracy. Dataset sizes and the choice of evaluation metrics greatly influence which method is considered successful.

## 5 Conclusions and Future Directions

Although adding LSDs as an auxiliary learning task significantly increases accuracy, it is unclear whether different shape descriptors could lead to further improvements. The LSDs proposed here were subjectively engineered based on features that we expected to be important to encode object shape. Future experiments could incorporate different features or focus on learning an optimal embedding rather than a hand-designed one. In that context, we note that it is not clear whether each component of the LSD embedding contributes equally to the improvement of affinity predictions.

Currently, we only use LSDs as an auxiliary learning task. As a result, affinities are still required to produce a segmentation. Whether this is really needed is an open question, since the predicted LSDs already identify objects reliably. An interesting future direction would be to use the predicted local shape information directly for fragment agglomeration. As an intermediate step, LSDs can serve to provide a second source of information for identifying errors in a segmentation. Once a segmentation is generated, LSDs could be calculated on the labels and then compared with the initial LSD predictions. The difference between the two would likely highlight regions containing errors (Figure 18.C).

LSDs were designed for the goal of neuron segmentation but might also be applicable to other instance segmentation problems. As an example, we found that the LSDs improve segmentations on plant epithelial cells (Figure 3, Table 3) and perform well on cell bodies and mitochondria (Figure 18.A,B). Since the LSDs are computed inside a Gaussian constrained to each object, the vectors allow for smooth transitions on both spherical and elongated shapes. In general, we believe that objects that have a blob-like structure such as other organelles and various cell types would likely benefit from LSDs. Furthermore, the direction vectors of the LSDs provide insight into neuropil vs. tract regions of the brain (Figure 18.D). These predictions could be leveraged in order to generate better tissue masks. While the LSDs presented here were conceived for a specific instance segmentation task, it would be interesting to see the LSDs extended and applied to other microscopy problems.

## 6 Acknowledgements

We thank Caroline Malin-Mayor, William Patton, and Julia Buhmann for code contributions; Nils Eckstein, Julia Buhmann and Steffen Wolf for helpful discussions; Stuart Berg for code to help with data acquisition; Jeremy Maitin-Shepard for helpful feedback on Neuroglancer; Viren Jain, Michał Januszewski, Jörgen Kornfeld, and Steve Plaza for access to data used for training and evaluation.

## Funding

This work was supported by the Howard Hughes Medical Institute. Uri Manor and Arlo Sheridan are supported by the Waitt Foundation, Core Grant application NCI CCSG (CA014195), NIH (R21 DC018237), NSF NeuroNex Award (2014862) and the Chan-Zuckerberg Initiative Imaging Scientist Award.

## Author Contributions

*Conceptualization*: Srinivas Turaga, Jan Funke. *Funding acquisition*: Stephan Saalfeld, Srinivas Turaga, Uri Manor, Jan Funke. *Software*: Arlo Sheridan, Diptodip Deb, Tri Nguyen, Jan Funke. *Data consolidation*: Arlo Sheridan, Diptodip Deb, Jan Funke. *Evaluation*: Arlo Sheridan, Jan Funke. *Data dissemination*: Arlo Sheridan, Jan Funke. *Visualization*: Arlo Sheridan, Jan Funke. *Writing (original draft)*: Arlo Sheridan, Jan Funke. *Writing (review and editing)*: Arlo Sheridan, Tri Nguyen, Diptodip Deb, Wei-Chung Allen Lee, Uri Manor, Jan Funke.

## Data Availability

All datasets analysed and/or generated during this study (raw data, training data, segmentations) are publicly available (see the “Data download” notebook on https://github.com/funkelab/lsd).

Figure 3: Zebrafinch: raw data, training data, segmentations

Figure 4: Hemi-brain: raw data, training data, segmentations

Figure 5: Fib-25: raw data, training data, segmentations

## Code Availability

The code used to train networks and segment neurons is available in the “LSD” repository, https://github.com/funkelab/lsd. Code used to evaluate the results is available in the “funlib.evaluate” repository, https://github.com/funkelab/funlib.evaluate.

## A Local Shape Descriptors

We define the notational shorthand

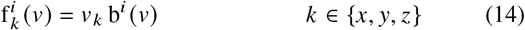

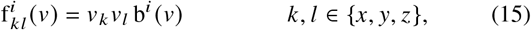

and use those to rewrite

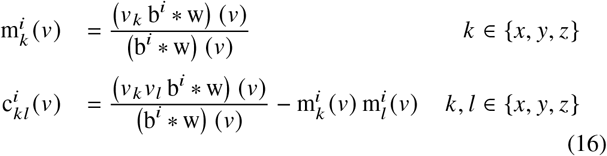

as follows:

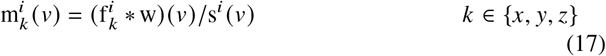

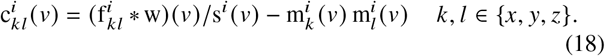

It can easily be seen that (17) is equal to the local center of mass, limited both by the object mask b and the local window w:

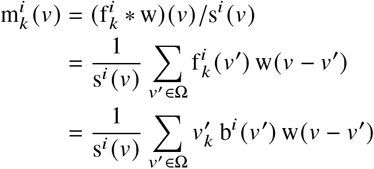

Similarly, the computation of the local covariance of voxel coordinates is equivalent to a convolution of the local window w with 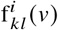. The local covariance is defined as:

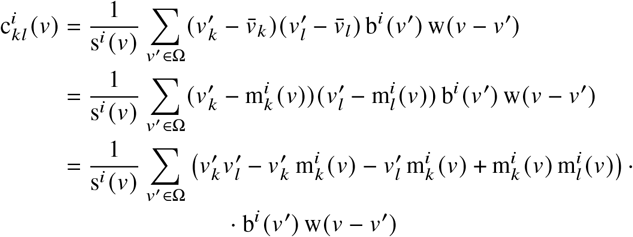

Rearranging terms reveals that 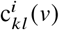 can efficiently be computed via a convolution as well:

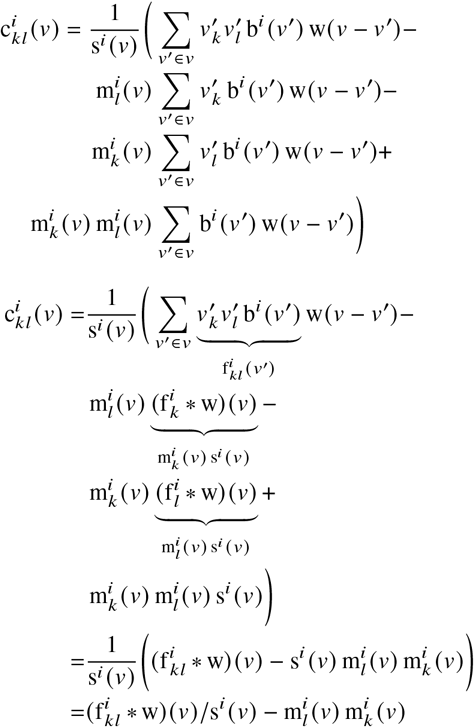

## B Min-Cut Metric

The Min-Cut Metric (MCM) measures the number of edit operations that need to be performed by a human annotator in a hypothetical proofreading tool that allows to: (1) *merge* wrongly split segments and (2) *split* wrongly merged segments by means of a min-cut^13^. To this end, we assume that the segmentation to interact with results from an agglomeration of fragments (or “supervoxels”). In particular, we assume that a *fragment graph G* = (*V, E, s*) is available, where each node *ν* ∈ *V* corresponds to a fragment and edges (*u, ν*) ∈ *E* are introduced between neighboring fragments. Each edge *e* ∈ *E* has an associated *merge score s*(*e*), which denotes under which agglomeration threshold the two incident fragments are to be merged into the same segment. A segmentation of the fragment graph is induced by a merge score threshold *θ*. Let *E* _*θ*_ = {*e* ∈ *E* | *s*(*e*) ≤ *θ*} be the set of *filtered edges*. Each connected component in the graph *G* _*θ*_ = (*V, E* _*θ*_) then corresponds to one segment. We will refer to the segment ID of a fragment under a given threshold *θ* as *l* _*θ*_(*ν*).

For the MCM metric, we assume ground-truth is available in the form of *skeletons*. Let *T* be the set of ground-truth skeletons, with each *t ∈ T* being a set of skeleton nodes. We will refer to the skeleton ID of a skeleton node *a* as *s a* and the fragment underlying the skeleton node as *l* _*θ*_ (*a*).

Given a segmentation *l* _*θ*_, MCM first simulates the splitting of all wrongly merged structures as given by ground-truth skeletons. For that, we first identify *merging* segments, *i.e*., segments that contain nodes from more than one skeleton. For each merging segment, we iteratively perform a series of min-cuts through the fragment graph, until the skeletons are separated. For that, we repeatedly find a pair of skeleton nodes *a* and *b*, such that

1. *s* (*a*) ≠ *s* (*b*) (the skeleton nodes belong to different skeletons),
2. *l* _*θ*_ (*a*) = *l* _*θ*_ (*b*) (the underlying fragments belong to the same segment), and
3. the Euclidean distance between *a* and *b* is minimized.

We then perform a min-cut on the fragment graph, with *u* and *ν* as the source and sink, respectively, and the capacity *c*(*e*) of edges *e* in the fragment graph set proportional to −*s*(*e*), such that edges with a high merge score are cheaper to cut. Once the mincut is found, all edges of the cut are removed from *E*_*θ*_ and the segmentation is updated accordingly. For a visualization of this procedure, see Supplementary Figure 17.A. This procedure aims to mimic a proofreader, who identified a merge and consequently picked two close locations on either side of the merge to perform a split operation.

In some cases, a min-cut can fail to separate all nodes of the merged skeletons with a single cut from two selected nodes. In this case, the procedure is repeated until all skeletons are separated, leading to additional split errors (see Supplementary Figure 17.B for an example).

After all skeletons are separated, the remaining split errors are counted. For that, we assume that each split requires one merge operation to be fixed. More generally, we identify the segments underlying each skeleton *t*: Let *L* (*t*) = {*l* _*θ*_ (*a*) | *a* ∈ *t*} be the set of segments underlying a skeleton. The number of required merge operations is then recorded as 1 − |*L* (*t*) |.

## C Zebrafinch

### C.1 Training

#### C.1.1 Data

33 volumes of densely labeled neurons^6^ were used for training. Each volume was padded with raw data. 30 volumes had raw dimensions of ∼ 7, 4.95, 4.95µm (zyx) and label dimensions of ∼ 3, 1.35, 1.35µm. The remaining 3 volumes had raw dimensions of ∼ 6.6, 5.9, 5.9µm and label dimensions of ∼ 2.6, 2.3, 2.3µm. Some regions containing glial processes were already set to zero and incorporated during network training (Supplementary Figure 12.A). A labels mask (1 inside labels RoI, null outside) was generated and used during training.

#### C.1.2 Networks

All methods used the U-Net architecture described in Funke et al. (2019). Networks consisted of three layers and were downsampled by a factor of [1,3,3] in the first two layers and [3,3,3] in the last layer. The reverse was done for the upsampling path. 12 initial feature maps were used and features were multiplied by a factor of 5 between layers. The resulting data was further convolved and passed through a sigmoid activation to get from 12 output feature maps^14^ to either 3 (affinities) or 10 feature maps (LSDs). All networks used an MSE loss, minimized with an Adam optimizer. The Malis network was trained to 10k iterations using MSE to initialize affinities and was then switched to Malis loss for the remainder of training.

Non auto-context networks had an input shape (raw) of [84,268,268] and output shape (labels, LSDs, affinities) of [48,56,56] (voxels, zyx). Auto-context networks had an input shape (raw) of [120,484,484], an intermediate shape (predicted LSDs) of [84,268,268], and an output shape (labels, affinities) of [48,56,56] (see Supplementary Figure 14.A for visualization of auto-context training shapes on Fib-25). The predicted LSDs used in the intermediate shape were taken from a pre-trained network which predicted LSDs from raw. Non auto-context networks were trained to 400k iterations. Auto-context networks were trained to ∼200k iterations following the 400k iterations of LSD training. See Supplementary Table 4 for a breakdown of the MtLsd network as an example.

All networks used a single voxel affinity neighborhood [1,1,1]. The LR network used three additional neighborhood steps of [3,3,3], [5,9,9] and [15,27,27]. The computed LSDs used a sigma of 120 nm and a downsampling factor of two.

#### C.1.3 Pipeline

Each training batch was randomly picked from one of the 33 training volumes. For each batch, the raw data was first normalized and padded with zeros. Labels were padded with the maximum padding required to contain at least 50% of ground-truth data assuming a worst case rotation of 45°. Data was randomly sampled from each dataset using a labels mask to ensure every batch contained at least 50% of ground-truth data. Data was then augmented with elastic transformations, random mirrors + transposes, and intensities (see Supplementary Table 4 for augmentation hyperparameters used for example MtLsd network). The following was done to the respective networks:

- **Baseline, LR** - Label boundaries were first eroded by a single voxel. Ground truth affinities were calculated on the labels using the pre-defined affinity neighborhoods and a scale array was created to balance loss between class labels. Training: [raw + gt affs] → pred affs.
- **Malis** - Label boundaries were eroded. If training loss was in Malis phase (*i.e*. after 10k iterations), connected components were relabelled before calculating ground-truth affinities. If training loss was in MSE phase (*i.e*. before 10k iterations), labels were subsequently balanced. Training: [raw + gt affs] → pred affs.
- **LSDs** -Ground truth LSDs were calculated on the labels using the pre-defined sigma and downsampling factor. Training: [raw + gt LSDs] → pred LSDs.
- **MtLsd** - Label boundaries were eroded. Ground truth LSDs were calculated followed by ground-truth affinities. Labels were then balanced. Training: [raw + gt LSDs + gt affs] → [pred LSDs + pred affs].
- **AcLsd, AcRLsd** - Label boundaries were eroded, ground-truth affinities were calculated, and labels were balanced. LSDs were then predicted in a slightly larger region and used as input to train the affinities. Training: [raw + gt LSDs] → pred LSDs → pred affs. For AcRLsd, cropped raw was incorporated in the second pass, in addition to predicted LSDs.

This process was repeated for a pre-defined number of iterations (generally until loss convergence).

### C.2 Prediction

Prediction was done in a block-wise fashion restricted to the Benchmark Roi. Individual workers used Gunpowder^4^ to predict output data (*i.e*. affinities or LSDs) and were distributed throughout the volume with DaisyNguyen et al. (2020). Block size was chosen with respect to how much data could fit in GPU memory. Most networks had a smaller block size in order to fit on 2080 RTX GPUs (∼ 12 GB RAM). While this increased the total number of blocks to process, the amount of workers available to use was sufficient to minimize total processing time. The auto-context networks were too large to fit on 2080 RTX GPUs and were therefore run on Tesla V100 GPUs. While there were less V100’s available, the block size could be greatly increased (∼ 32 GB RAM), decreasing the total amount of blocks to process. LSDs were physically written to file before use in the auto-context networks. This was done for visualization; prediction could be adapted to generate LSDs on the fly. All networks wrote affinities to file and then subsequently used them for segmentation.

### C.3 Segmentation

#### C.3.1 Watershed

Seeded watershed^15^ was done on the affinities generated during prediction. Both non-masked and neuropil-masked supervoxels were produced. Due to data anisotropy, supervoxels were extracted for each section separately. An epsilon agglomeration was used to agglomerate fragments to a predefined threshold (0.1). This was done to decrease the number of RAG nodes during the full agglomeration step. Supervoxels which had an average affinity value lower than a pre-defined threshold (0.05) were filtered out of the RAG and set to zero in the resulting datasets. A block size of 3.6µm^3^ and context of [12,27,27] voxels (zyx) were used.

#### C.3.2 Agglomeration

Supervoxels were agglomerated using hierarchical region agglomeration^15^ in which edges with lower affinity scores are merged earlier. We empirically chose to use both 50 and 75 quantile merge functions since they produced the best results in Funke et al. (2019). The same block size/context as watershed were used.

#### C.3.3 Segment

The center point of the Benchmark Roi was used to grow sub RoIs. The first sub RoI was created by growing the center point in each dimension (positive and negative) by the block size used during watershed and agglomeration. This resulted in 10.8µm edge lengths (3.6µm + (3.6 × 2)). This RoI was again grown by the block size to produce an RoI with 18µm edge lengths (10.8µm + (3.6 × 2)). This was repeated for a total of 10 RoIs (in addition to the Benchmark Roi). Segmentations were created for each RoI by cropping the RAG and relabelling connected components on the graph^16^. This was done over a range of thresholds for each network (threshold range = [0 - 1], step size = 0.02, total thresholds = 50). Segmentations were created for both masked / non-masked data and both merge functions.

### C.4 Evaluation

Manually traced skeletons^6^ (12 validation, 50 testing) were used for evaluation. For each sub RoI, skeletons were cropped, either masked to neuropil or not masked, and connected components were relabelled^16^. For affinity-based methods, fragment ids were first mapped to skeleton ids in each block. This mapping was then used to assign segment ids to skeleton ids for each threshold, using the lookup tables generated in Supplementary Section C.3.3. A site mask was used to restrict segments to the skeleton nodes. The resulting node - segment mapping was used to compute^17^ ERL, NERL and VoI.

Additionally, on the first three sub RoIs, the MCM was calculated using masked skeletons and segmentations. For the FFN, a single segmentation^6^ was used to generate the node - segment mappings. The full segmentation was downloaded^18^ and cropped to each sub RoI, either masked or not masked, and connected components were relabelled. Only ERL, NERL and VoI were calculated as there were no supervoxels to use for the MCM. For all affinity-based methods, we repeated these steps using the validation skeletons on the benchmark RoI. The optimal thresholds indicated in test set plots were determined by the thresholds which minimized VoI (for VoI and MCM plots) and maximized ERL (for ERL plots).

## D FIB-SEM volumes

### D.1 Training

#### D.1.1 Data

**H****emi-brain**: 8 volumes of densely labeled neurons^7^ were used for training. Volumes were taken from various neuropils^19^ contained within the dataset generated in (Scheffer et al., 2020). The Lobula Plate and Lateral Horn volumes contained 4µm^3^ of raw data and ~2µm^3^ of labeled data, while the others contained ~6µm^3^ of raw data and ~4µm^3^ of labeled data (Supplementary Figure 12.B).

**Fib-25**: 4 volumes of densely labeled neurons^7^ were used for training. The labels were not padded with raw as was done in the Zebrafinch and Hemi-brain volumes. Two volumes contained RoI 4µm^3^ of raw / labeled data, and two volumes contained 2µm^3^ of raw / labeled data (Supplementary Figure 12.C). Label masks were generated for all volumes, as done in the Zebrafinch.

#### D.1.2 Networks

Networks consisted of same architecture as Zebrafinch networks except downsampling was isotropic with a factor of [2,2,2] in the first two layers and [3,3,3] in the last layer. Features were multiplied by a factor of 6 between layers.

Non auto-context networks had an input shape (raw) of [196,196,196] and output shape (labels, LSDs, affinities) of [92,92,92]. Auto-context networks had an input shape (raw) of [304,304,304], an intermediate shape (predicted LSDs) of [196,196,196], and an output shape (labels, affinities) of [92,92,92]. Non auto-context networks were trained to 400k iterations. Auto-context networks were trained to ∼ 300k iterations following 400k iterations of LSD training. See Supplementary Table 5 for a breakdown of the MtLsd network as an example.

All networks used a single voxel affinity neighborhood [1,1,1]. The LR network used three additional neighborhood steps of [3,3,3], [5,5,5] and [13,13,13]. The computed LSDs used a sigma of 80 nm and a downsampling factor of two.

#### D.1.3 Pipeline

All networks were trained following the same pipeline as the Zebrafinch networks using either 8 (Hemi-brain) or 4 (Fib-25) ground-truth volumes. The augmentations were computed isotropically, in contrast to the Zebrafinch networks (see Supplementary Table 5 for augmentation hyper-parameters used for example MtLsd network). For Fib-25 training, affinities and LSDs were masked at the boundaries to ensure that prediction on the irregularly shaped Fib-25 volume did not include boundary artifacts (Supplementary Figure 14.B).

### D.2 Prediction

For the Hemi-brain, prediction was restricted to the three RoIs described in Section 3.3.2. For Fib-25, prediction was done on the full Fib-25 volume (including the background). The process was the same as in the Zebrafinch.

### D.3 Segmentation

#### D.3.1 Watershed

Fragment extraction was performed isotropically and used no epsilon agglomeration step or mean affinity filtering, in contrast to the Zebrafinch. A block size of 3µm^3^ and context of 31 voxels were used. For the Hemi-brain, watershed was done on each predicted RoI and restricted using an Ellipsoid Body mask^7^. For Fib-25, an irregularly shaped tissue mask^6^ was used^20^.

#### D.3.2 Agglomeration

Agglomeration was done using the same merge functions on the Zebrafinch. The same block size and context from watershed were used.

#### D.3.3 Segment

For the Hemi-brain, segmentations were created for the three processed RoIs. For Fib-25, segmentations were created for the full Fib-25 RoI and two sub RoIs. The same threshold range from the Zebrafinch was used.

### D.4 Evaluation

**H****emi-brain**: dense ground-truth^7^ and an FFN segmentation^6^ were available for the entire Hemi-brain. Both volumes were downloaded^21^,^22^ and cropped to the three established RoIs. The cropped datasets were then constrained to the Ellipsoid Body. The ground-truth was filtered using a whitelist of proofread ids. Connected components were relabelled and boundaries were slightly eroded. **Fib-25**: dense ground-truth^7^ and an FFN segmentation^6^ were already cropped to the testing RoI. The ground-truth was already filtered with a whitelist and boundaries were already eroded. Both volumes were further cropped to the two sub RoIs and connected components were relabelled. For affinity-based methods, VoI was calculated between the consolidated ground-truth and segmentations over all thresholds on each RoI. For the FFN, VoI was calculated on the single segmentation for each RoI.

## E Throughput

For each affinity-based network, we calculated the amount of floating point operations (FLOPS) for the processing of one block^23^ using TensorFlow’s Profiler^24^ (see Supplementary Table 6 for a breakdown by operation). From the computed FLOPS and the block size, we derived FLOPS/ µm.

For FFN, FLOPS are reported for the full Zebrafinch RoI in Januszewski et al. (2018), which we divided by the full RoI volume to get FLOPS/µm.

## F Extended Experiments

### F.1 Serial Section Data (ssTEM)

To test the efficacy of LSDs on both ssTEM and mouse neural tissue, we used the publicly available data from (Microns Consortium et al., 2021). To train the networks, we used 38 volumes (∼ 908µm^3^ total, ∼ 24µm^3^ average) taken from three datasets (Basil, Minnie, Pinky). Several volumes were excluded from training and testing due to overlap. The volumes were either imaged at 40×4×4 or 40×8×8 nanometer resolution (zyx). We therefore chose to downsample volumes by a factor of 2 laterally to ensure consistency. Networks were trained similarly to the Zebrafinch networks with added missing and shifted section augmentations.

Due to the anisotropy of the data and valid padding of the networks, we opted to use 2D convolutions in the lowest level of the U-NET. We trained Baseline, LR, MtLsd, AcLsd, and AcRLsd networks and omitted Malis following the results on the block face datasets. A foreground mask was provided (zero in missing or broken sections) and used to ensure that predictions persisted through these regions.

Evaluation was done on 4 test volumes (∼232µm^3^ total, ∼58µm^3^ average). Watershed and agglomeration was done similarly to the Zebrafinch (with the exception of a neuropil mask). Ground truth neurons were relabelled between missing sections to gauge neuropil performance since we did not engineer ssTEM-specific preprocessing (improved alignment, duplicated section augmentations, etc.)

Our results suggest that the LSDs also work well on serial section data and would likely benefit from the same processing as other methods. They look qualitatively reasonable (Figure 1) and when considering pure neuropil, they seem to outperform baseline methods (Table 1). Consistent with the other datasets, using an autocontext network produces the highest accuracy (AcLsd VoI of 0.639). Due to the small size of these datasets, it is difficult to extrapolate accuracy to larger volumes but we expect that the LSDs would generally help to improve Baseline affinities.

### F.2 Ablations

In order to see how important each component of the LSDs is, we ran an ablation experiment in which we limited the embedding to each possible combination of components for both an MtLsd network and AcLsd network. Networks were trained on the Hemi-brain training data and tested on the 12 µm RoI.

We expected that the MtLsd networks would not be significantly affected by removing components since they still learn the affinities in the same pass. We found that to generally be true, with the results not indicative of a single superior and inferior combination (Table 2). Conversely, we hypothesized that the AcLsd networks would be highly dependent on which components of the LSDs were fed into the second pass. We found that, generally speaking, any combination of LSDs will generate reasonable results but using only the size component as input to the second pass produces poor results (Table 2). This is illustrated in Figure 2, where it is clear that the affinities and subsequent segmentations do benefit from a combination of components that is not limited to the size of the neural process. It is also not clear how much the numbers are due to random training noise.

### F.3 Non-EM Data: Epithelial Cells

We tested the LSDs on the publicly available plant epithelial ovule dataset from Wolny et al. (2020). We used 20 volumes for training (∼ 6706µm^3^ total, ∼ 335µm^3^ average). All volumes were cropped in the z dimension to sections that contained labels (i.e, volume ends that were completely background were removed). We implemented networks as done for the ssTEM extended experiment, but found results to be poor when using hierarchical agglomeration due to the challenging nature of the data. To overcome this, we decided to use the mutex watershed (Wolf et al., 2018). To this end, we did not consider Baseline affinities since a long range affinity neighborhood is required for the mutex watershed. Additionally, we found MtLsd results using a long range affinity neighborhood to be qualitatively poor, likely due to the network trying to learn too much simultaneously.

**Supplementary Table 1:** Best Variation of Information (Sum) per network on four testing volumes from microns data (lower is better). Best method per volume is shown in bold.

| Volume | BASELINE | LR | MtLSD | AcLSD | AcRLSD |
| --- | --- | --- | --- | --- | --- |
| basil_v001 | 0.235 | 0.206 | **0.186** | 0.210 | 0.445 |
| basil_v004 | 0.775 | 0.789 | 0.830 | **0.658** | 0.979 |
| minnie_v024 | 1.594 | 1.978 | 1.823 | **1.440** | 1.963 |
| pinky_v104 | 0.244 | **0.229** | 0.2355 | 0.248 | 0.245 |
| Avg. ± SD | 0.712 ± 0.554 | 0.800 ± 0.719 | 0.769 ± 0.659 | **0.639 ± 0.495** | 0.908 ± 0.666 |

**Supplementary Figure 1:**
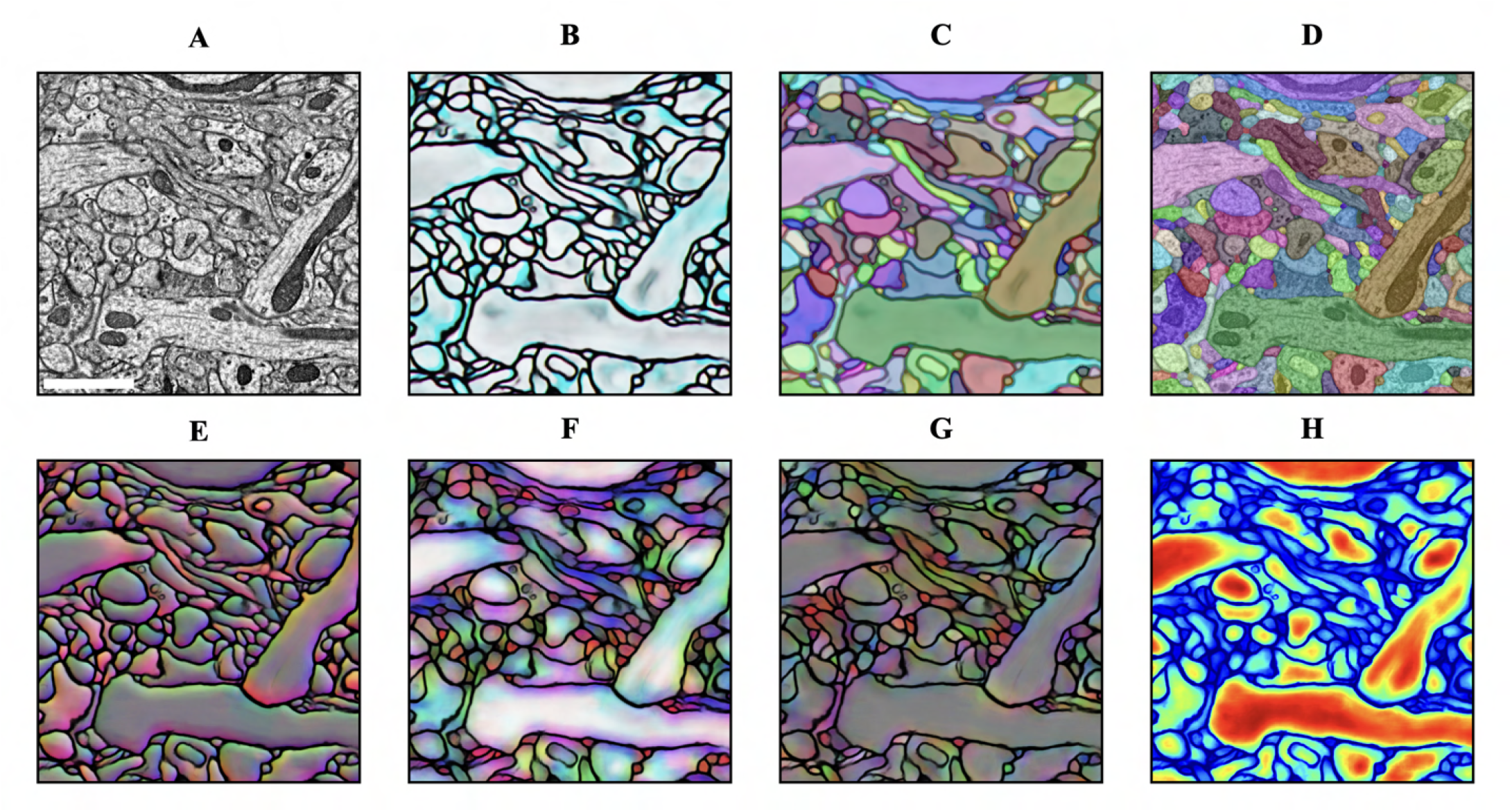
LSD results on example ssTEM data. **A**. Raw EM data, scale bar = 1 µm. **B**. Affinities computed from second pass of AcLsd network. **C**. Segmentation produced from affinities. **D**. Segmentation overlayed with raw data. **E**,**F**,**G**,**H**. Mean offset, diagonal entries of covariance, off-diagonal entries of covariance, and size components of LSDs.

We therefore limited our evaluation to LR affinities as a baseline, and an AcLsdnetwork. In contrast to the Zebrafinch, Fib-25, and Hemi-brain experiments, our long range neighborhood also consisted of diagonal affinities as they are useful for the mutex watershed. We used 7 volumes for testing (∼ 2466µm^3^ total, ∼ 352µm^3^ average). Following the mutex watershed, we filtered out small objects (500 voxels minimum size) and expanded the labels to fill the intermediate space between objects. Since some of the ground truth was not fully labeled, we restricted our segmentations to a foreground mask generated from the ground truth. Finally, we relabelled connected components. Results suggest that the LSDs are useful for non-EM data, and seem to work especially well when combined with the mutex watershed (Table 3), a promising indication for future work. The results are also qualitatively appealing (Figure 3).

**Supplementary Table 2:** Best Variation of Information (Sum) per component combination for both networks (lower is better).

| Component | MtLSD | AcLSD |
| --- | --- | --- |
| Off-Diagonals of Covariance | 0.1295 | 0.0268 |
| Off-Diagonals of Covariance + Size | 0.0332 | 0.0613 |
| Full LSDs | 0.0373 | 0.0261 |
| Mean Offsets | 0.0406 | 0.0344 |
| Mean Offsets + Off-Diagonals of Covariance | 0.0727 | 0.0457 |
| Mean Offsets + Off-Diagonals of Covariance + Size | 0.1681 | 0.0439 |
| Mean Offsets + Diagonals of Covariance | 0.0412 | 0.0299 |
| Mean Offsets + Diagonals of Covariance + Off-Diagonals of Covariance | 0.0465 | **0.0256** |
| Mean Offsets + Diagonals of Covariance + Size | **0.0258** | 0.0294 |
| Mean Offsets + Size | 0.0835 | 0.0749 |
| Diagonals of Covariance | 0.0491 | 0.0329 |
| Diagonals of Covariance + Off-Diagonals of Covariance | 0.0374 | 0.1168 |
| Diagonals of Covariance + Off-Diagonals of Covariance + Size | 0.0414 | 0.0421 |
| Diagonals of Covariance + Size | 0.0758 | 0.0295 |
| Size | 0.0463 | 0.8444 |

**Supplementary Figure 2:**
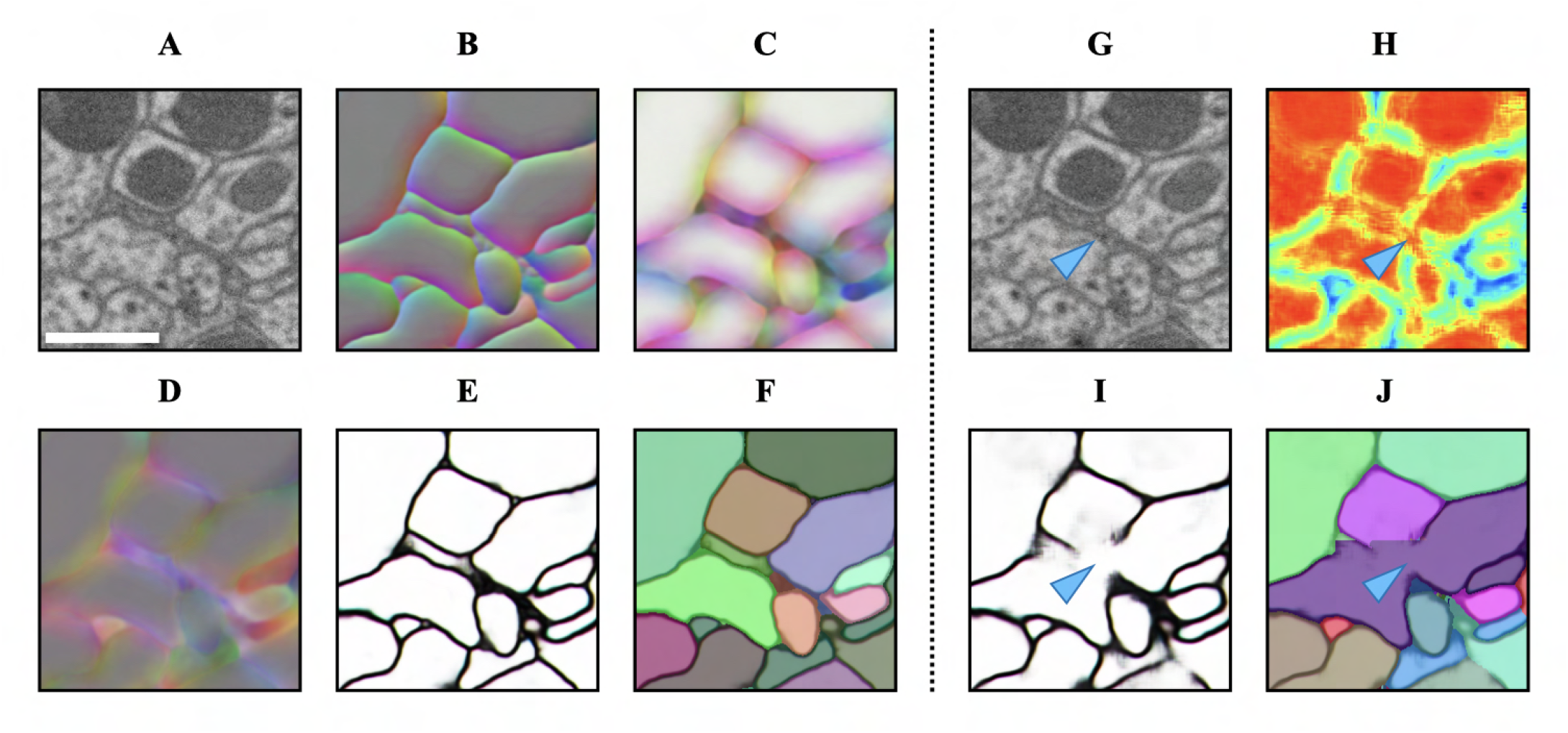
Results from best autocontext ablation components (left of dashed line) and worst components (right of dashed line) on Hemi-brain cutout. **A, G**. Raw EM data, scale bar = 500 nm. **B**,**C**,**D**. Mean offset, orthogonals, and diagonals of LSDs in best ablation. Best performing ablation did not include a size component. **E, F**. Affinities and segmentation produced from best ablation. **H**. Size (and sole) component of LSDs in worst ablation. **I, J**. Affinities and segmentation produced from worst ablation. Autocontext using just the size component as input causes large gaps in the affinities resulting in a false merge (blue arrows).

**Supplementary Table 3:** Best Variation of Information (Sum) for each network over seven testing volumes (lower is better).

| Volume | LR | AcLSD |
| --- | --- | --- |
| N_522 | 1.881 | **1.388** |
| N_593 | **1.644** | 1.740 |
| N_441 | 1.462 | **1.196** |
| N_435 | 1.671 | **1.495** |
| N_590 | **1.474** | 1.636 |
| N_511 | **1.153** | 1.290 |
| N_294 | 2.021 | **1.604** |
| Avg. ± SD | 1.615 ± 0.266 | **1.478 ± 0.182** |

**Supplementary Figure 3:**
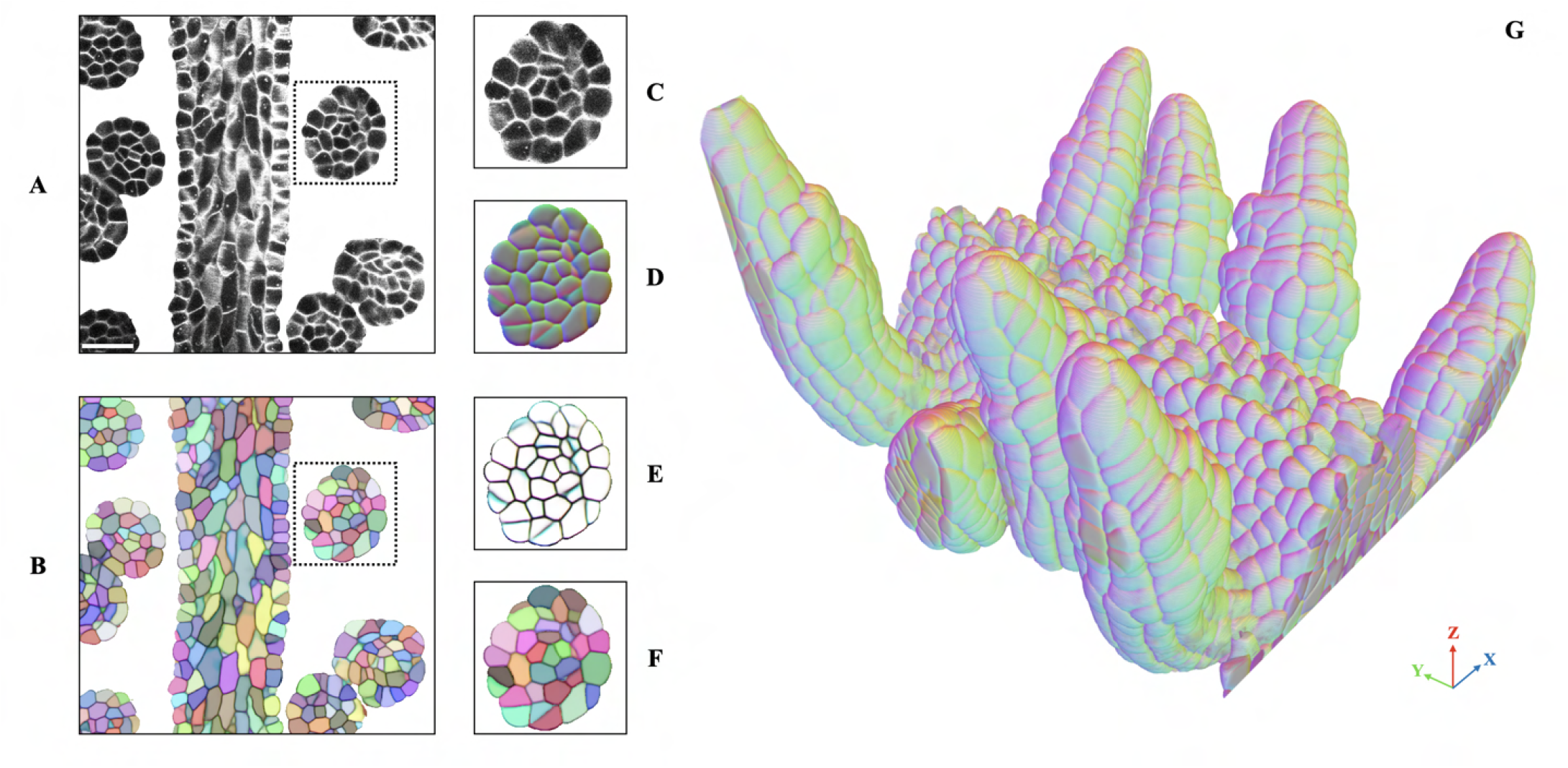
LSDs on plant epithelial cell data. **A**. Raw microscopy data from example volume, scale bar = 1 µm. **B**. Segmented cells. **C, D, E, F**. Insets showing raw data, mean offset component of LSDs, resulting affinities and segmentation from second pass of AcLsd network. **G**. 3D rendering of the mean offset component of LSDs. Cartesian coordinate inset shows corresponding directions.

**Supplementary Table 4:** Training parameters and augmentations^4^ of **MtLsd** network on Zebrafinch dataset.

| Parameter | Value |
| --- | --- |
| Input feature maps | 12 |
| Layer fmap scale | 5 |
| Downsampling factors | [[1,3,3],[1,3,3],[3,3,3]] |
| Output feature maps | 14 |
| Input shape | [84,268,268] |
| Output shape | [48,56,56] |
| Loss | MSE |
| Optimizer | Adam |
| Learning rate | $0.5 \times 10^{-4}$ |
| $\beta_1$ | 0.95 |
| $\beta_2$ | 0.999 |
| $\epsilon$ | $1 \times 10^{-8}$ |
| Iterations | 400,000 |

| Augmentation | Parameter | Value |
| --- | --- | --- |
| Elastic | control point spacing | (4,4,10) |
|  | jitter sigma | (0, 2, 2) |
|  | subsample | 8 |
| Rotation | axis | x,y,z |
| | angle | in $[0, 2\pi]$ |
| Section Defects | slip probability | 0.05 |
|  | shift probability | 0.05 |
|  | max misalign | 10 |
| Mirror | axes | x,y,z |
| Transpose | axes | x, y |
| Intensity | scale | in $[0.9, 1.1]$ |
| | shift | in $[-0.1, 0.1]$ |

**Supplementary Figure 4:**
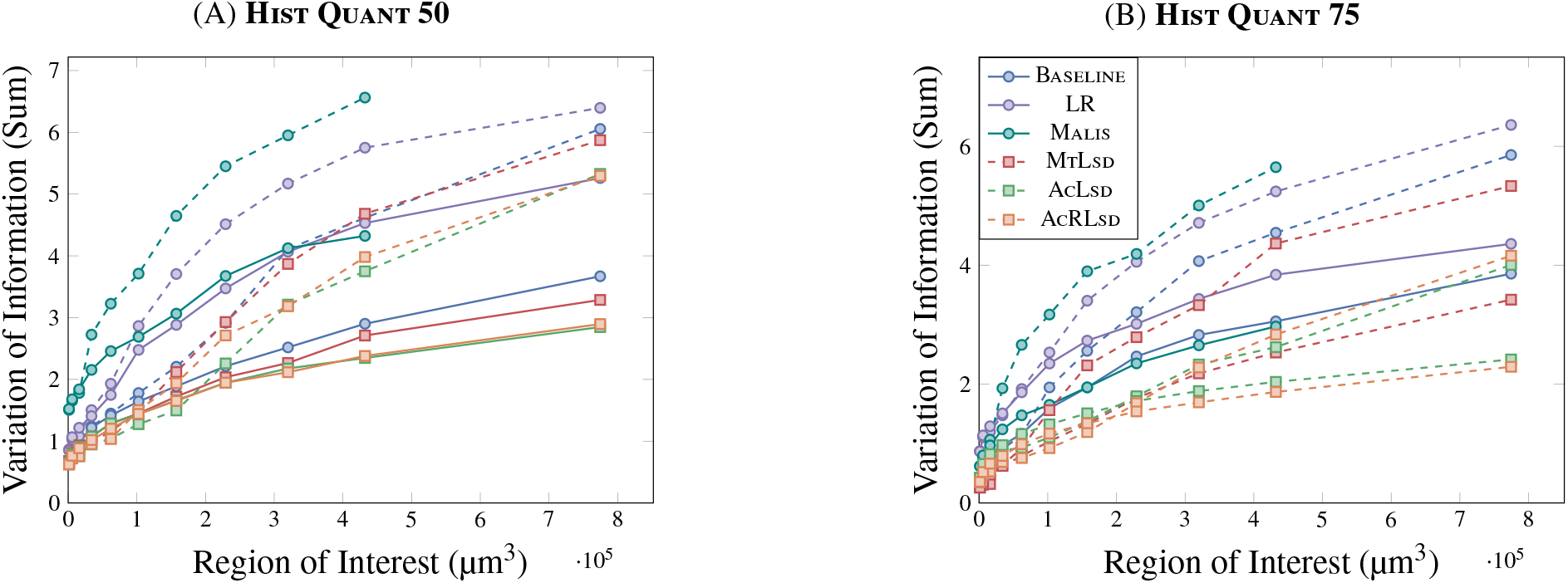
VoI Sum vs RoI on masked (solid) and non-masked (dashed) Zebrafinch data.

**Supplementary Table 5:** Training parameters and augmentations^4^ of **MtLsd** network on FIB-SEM datasets.

| Parameter | Value |
| --- | --- |
| Input feature maps | 12 |
| Layer fmap scale | 6 |
| Downsampling factors | [[2,2,2],[2,2,2],[3,3,3]] |
| Output feature maps | 14 |
| Input shape | [196,196,196] |
| Output shape | [92,92,92] |
| Loss | MSE |
| Optimizer | Adam |
| Learning rate | $0.5 \times 10^{-4}$ |
| $\beta_1$ | 0.95 |
| $\beta_2$ | 0.999 |
| $\epsilon$ | $1 \times 10^{-8}$ |
| Iterations | 400,000 |

| Augmentation | Parameter | Value |
| --- | --- | --- |
| Elastic 1 | control point spacing | (40,40,40) |
|  | jitter sigma | (0, 0, 0) |
|  | subsample | 8 |
| Rotation 1 | axis | x,y,z |
| | angle | in $[0, 2\pi]$ |
| Section Defects 1 | slip probability | 0 |
|  | shift probability | 0 |
|  | max misalign | 0 |
| Mirror | axes | x,y,z |
| Transpose | axes | x,y,z |
| Elastic 2 | control point spacing | (40,40,40) |
|  | jitter sigma | (2, 2, 2) |
|  | subsample | 8 |
| Rotation 2 | axis | x,y,z |
| | angle | in $[0, 2\pi]$ |
| Section Defects 2 | slip probability | 0.01 |
|  | shift probability | 0.01 |
|  | max misalign | 1 |
| Intensity | scale | in $[0.9, 1.1]$ |
| | shift | in $[-0.1, 0.1]$ |

**Supplementary Figure 5:**
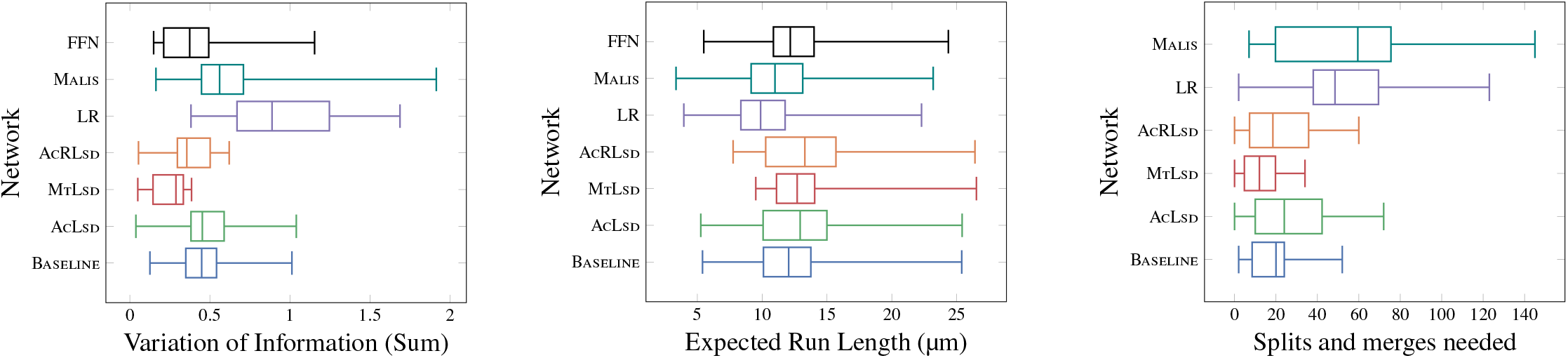
Metric distribution across 11 randomly sampled non-masked sub RoIs in the Zebrafinch.

**Supplementary Table 6:** FLOPs breakdown by operation for **MtLsd** on Zebrafinch dataset.

| Operation | FLOPs |
| --- | --- |
| unet_layer_2_left_1/convolution | 940584960000 |
| unet_layer_2_right_0/convolution | 786542400000 |
| unet_layer_3_left_1/convolution | 419904000000 |
| unet_layer_1_left_1/convolution | 416320819200 |
| unet_layer_2_right_1/convolution | 338411520000 |
| unet_layer_1_right_0/convolution | 226738828800 |
| unet_layer_2_left_0/convolution | 208980000000 |
| unet_layer_0_left_1/convolution | 164826316800 |
| unet_layer_3_left_0/convolution | 123832800000 |
| unet_layer_1_right_1/convolution | 105305702400 |
| unet_layer_1_left_0/convolution | 87354028800 |
| unet_layer_0_right_0/convolution | 84455094528 |
| gradients/unet_up_3_to_2/conv3d_transpose_grad/Conv3D | 83980800000 |
| unet_layer_0_right_1/convolution | 46982799360 |
| gradients/unet_up_2_to_1/conv3d_transpose_grad/Conv3D | 22560768000 |
| unet_layer_0_left_0/convolution | 14151319488 |
| gradients/unet_up_1_to_0/conv3d_transpose_grad/Conv3D | 7020380160 |
| embedding_0/convolution | 1242931200 |
| affs_0/convolution | 372879360 |
| unet_layer_0_left_0/BiasAdd | 262061472 |
| gradients/unet_layer_0_left_0/BiasAdd_grad/BiasAddGrad | 262061460 |
| gradients/AddN_3 | 254361600 |
| unet_layer_0_left_1/BiasAdd | 254361600 |
| gradients/unet_layer_0_left_1/BiasAdd_grad/BiasAddGrad | 254361588 |
| unet_layer_1_left_0/BiasAdd | 134805600 |
| gradients/unet_layer_1_left_0/BiasAdd_grad/BiasAddGrad | 134805540 |
| unet_layer_1_left_1/BiasAdd | 128494080 |
| gradients/AddN_2 | 128494080 |
| gradients/unet_layer_1_left_1/BiasAdd_grad/BiasAddGrad | 128494020 |
| unet_layer_3_left_1/kernel/Initializer/random_uniform | 60750000 |
| mean_squared_error/num_present | 44390399 |
| mean_squared_error/SquaredDifference | 88780800 |
| unet_layer_0_right_0/BiasAdd | 65165968 |
| gradients/unet_layer_0_right_0/BiasAdd_grad/BiasAddGrad | 65165954 |
| unet_layer_2_left_0/BiasAdd | 64500000 |
| gradients/unet_layer_2_left_0/BiasAdd_grad/BiasAddGrad | 64499700 |
| unet_layer_0_right_1/BiasAdd | 62146560 |
| gradients/AddN | 62146560 |
| gradients/unet_layer_0_right_1/BiasAdd_grad/BiasAddGrad | 62146546 |
| unet_up_1_to_0/BiasAdd | 58503168 |
| gradients/unet_up_1_to_0/BiasAdd_grad/BiasAddGrad | 58503156 |
| unet_layer_2_left_1/BiasAdd | 58060800 |
| gradients/AddN_1 | 58060800 |
| gradients/unet_layer_2_left_1/BiasAdd_grad/BiasAddGrad | 58060500 |
| embedding_0/BiasAdd | 44390400 |
| gradients/mean_squared_error/Mul_grad/mul | 44390400 |
| gradients/mean_squared_error/Mul_grad/mul_1 | 44390400 |
| gradients/mean_squared_error/SquaredDifference_grad/Neg | 44390400 |
| gradients/mean_squared_error/SquaredDifference_grad/mul | 44390400 |
| mean_squared_error/Mul | 44390400 |
| gradients/mean_squared_error/SquaredDifference_grad/mul_1 | 44390400 |
| gradients/mean_squared_error/SquaredDifference_grad/sub | 44390400 |
| mean_squared_error/Sum | 44390399 |
| gradients/embedding_0/BiasAdd_grad/BiasAddGrad | 44390390 |
| gradients/mean_squared_error/Mul_grad/Sum_1 | 39951360 |
| unet_up_2_to_1/BiasAdd | 37601280 |
| gradients/unet_up_2_to_1/BiasAdd_grad/BiasAddGrad | 37601220 |
| unet_layer_1_right_0/BiasAdd | 34990560 |
| gradients/unet_layer_1_right_0/BiasAdd_grad/BiasAddGrad | 34990500 |
| unet_layer_1_right_1/BiasAdd | 32501760 |
| gradients/unet_layer_1_right_1/BiasAdd_grad/BiasAddGrad | 32501700 |
| unet_up_3_to_2/BiasAdd | 27993600 |
| gradients/unet_up_3_to_2/BiasAdd_grad/BiasAddGrad | 27993300 |
| mean_squared_error_1/SquaredDifference | 26634240 |
| mean_squared_error_1/num_present | 13317119 |
| unet_layer_3_left_0/kernel/Initializer/random_uniform | 12150000 |
| unet_up_3_to_2/kernel/Initializer/random_uniform | 12150000 |
| unet_layer_2_right_0/BiasAdd | 24276000 |
| gradients/unet_layer_2_right_0/BiasAdd_grad/BiasAddGrad | 24275700 |
| unet_layer_2_right_1/BiasAdd | 20889600 |
| gradients/unet_layer_2_right_1/BiasAdd_grad/BiasAddGrad | 20889300 |
| gradients/mean_squared_error_1/SquaredDifference_grad/mul | 13317120 |
| gradients/mean_squared_error_1/SquaredDifference_grad/Neg | 13317120 |
| gradients/mean_squared_error_1/SquaredDifference_grad/mul_1 | 13317120 |
| mean_squared_error_1/Mul | 13317120 |
| gradients/mean_squared_error_1/Mul_grad/mul_1 | 13317120 |
| gradients/mean_squared_error_1/Mul_grad/mul | 13317120 |
| affs_0/BiasAdd | 13317120 |
| gradients/mean_squared_error_1/SquaredDifference_grad/sub | 13317120 |
| mean_squared_error_1/Sum | 13317119 |
| gradients/affs_0/BiasAdd_grad/BiasAddGrad | 13317117 |
| unet_layer_2_right_0/kernel/Initializer/random_uniform | 4860000 |
| unet_layer_3_left_0/BiasAdd | 7644000 |
| gradients/unet_layer_3_left_0/BiasAdd_grad/BiasAddGrad | 7642500 |
| unet_layer_3_left_1/BiasAdd | 5184000 |
| gradients/unet_layer_3_left_1/BiasAdd_grad/BiasAddGrad | 5182500 |
| unet_layer_2_right_1/kernel/Initializer/random_uniform | 2430000 |
| unet_layer_2_left_1/kernel/Initializer/random_uniform | 2430000 |
| unet_layer_2_left_0/kernel/Initializer/random_uniform | 486000 |
| unet_layer_1_right_0/kernel/Initializer/random_uniform | 194400 |
| unet_up_2_to_1/kernel/Initializer/random_uniform | 162000 |
| unet_layer_1_right_1/kernel/Initializer/random_uniform | 97200 |
| unet_layer_1_left_1/kernel/Initializer/random_uniform | 97200 |
| unet_layer_1_left_0/kernel/Initializer/random_uniform | 19440 |
| unet_layer_0_right_0/kernel/Initializer/random_uniform | 9072 |
| unet_up_1_to_0/kernel/Initializer/random_uniform | 6480 |
| unet_layer_0_right_1/kernel/Initializer/random_uniform | 5292 |
| unet_layer_0_left_1/kernel/Initializer/random_uniform | 3888 |
| unet_layer_0_left_0/kernel/Initializer/random_uniform | 324 |
| embedding_0/kernel/Initializer/random_uniform | 140 |
| affs_0/kernel/Initializer/random_uniform | 42 |
| gradients/Slice_1_grad/sub | 5 |
| gradients/Slice_1_grad/sub_1 | 5 |
| gradients/Slice_2_grad/sub | 5 |
| gradients/Slice_2_grad/sub_1 | 5 |
| gradients/Slice_grad/sub | 5 |
| gradients/Slice_grad/sub_1 | 5 |
| mean_squared_error_1/Greater | 1 |
| unet_up_1_to_0/mul | 1 |
| unet_up_3_to_2/mul_2 | 1 |
| unet_up_3_to_2/mul_1 | 1 |
| unet_up_3_to_2/mul | 1 |
| unet_up_3_to_2/add_2 | 1 |
| unet_up_3_to_2/add_1 | 1 |
| unet_up_3_to_2/add | 1 |
| unet_up_2_to_1/mul_2 | 1 |
| unet_up_2_to_1/mul_1 | 1 |
| unet_up_2_to_1/mul | 1 |
| unet_up_2_to_1/add_2 | 1 |
| unet_up_2_to_1/add_1 | 1 |
| unet_up_2_to_1/add | 1 |
| unet_up_1_to_0/mul_2 | 1 |
| unet_up_1_to_0/mul_1 | 1 |
| gradients/mean_squared_error_1/div_grad/RealDiv_2 | 1 |
| Adam/mul_1 | 1 |
| add | 1 |
| gradients/mean_squared_error/div_grad/Neg | 1 |
| gradients/mean_squared_error/div_grad/RealDiv | 1 |
| gradients/mean_squared_error/div_grad/RealDiv_1 | 1 |
| gradients/mean_squared_error/div_grad/RealDiv_2 | 1 |
| gradients/mean_squared_error/div_grad/mul | 1 |
| gradients/mean_squared_error_1/div_grad/Neg | 1 |
| gradients/mean_squared_error_1/div_grad/RealDiv | 1 |
| gradients/mean_squared_error_1/div_grad/RealDiv_1 | 1 |
| unet_up_1_to_0/add_2 | 1 |
| gradients/mean_squared_error_1/div_grad/mul | 1 |
| mean_squared_error/Equal | 1 |
| mean_squared_error/Greater | 1 |
| mean_squared_error/div | 1 |
| mean_squared_error_1/Equal | 1 |
| Adam/mul | 1 |
| mean_squared_error_1/div | 1 |
| unet_up_1_to_0/add | 1 |
| unet_up_1_to_0/add_1 | 1 |
| Total | 4083673765073 |

**Supplementary Figure 6:**
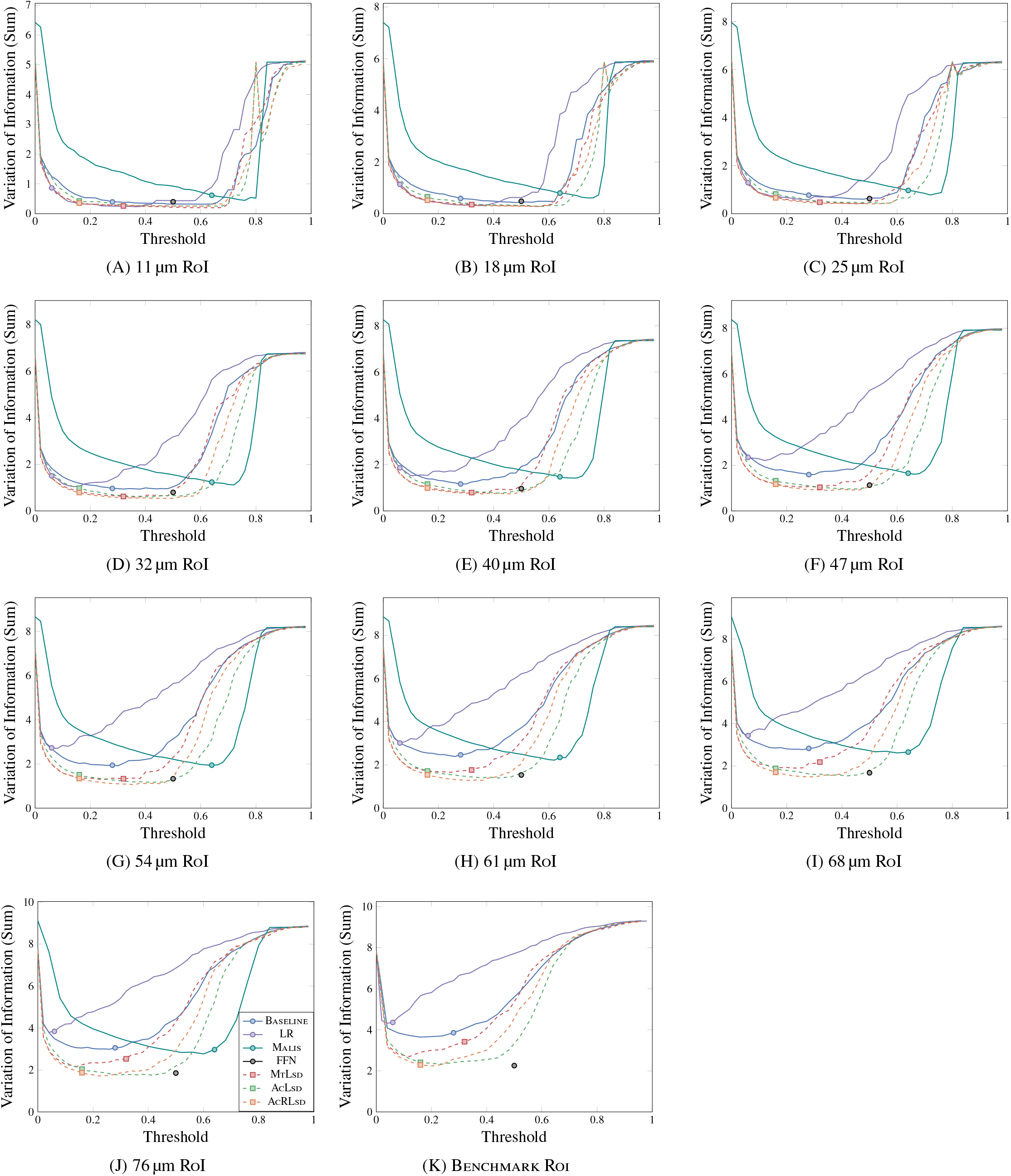
VoI Sum vs threshold across Zebrafinch RoIs. Points correspond to thresholds which minimized VoI Sum on the validation dataset.

**Supplementary Figure 7:**
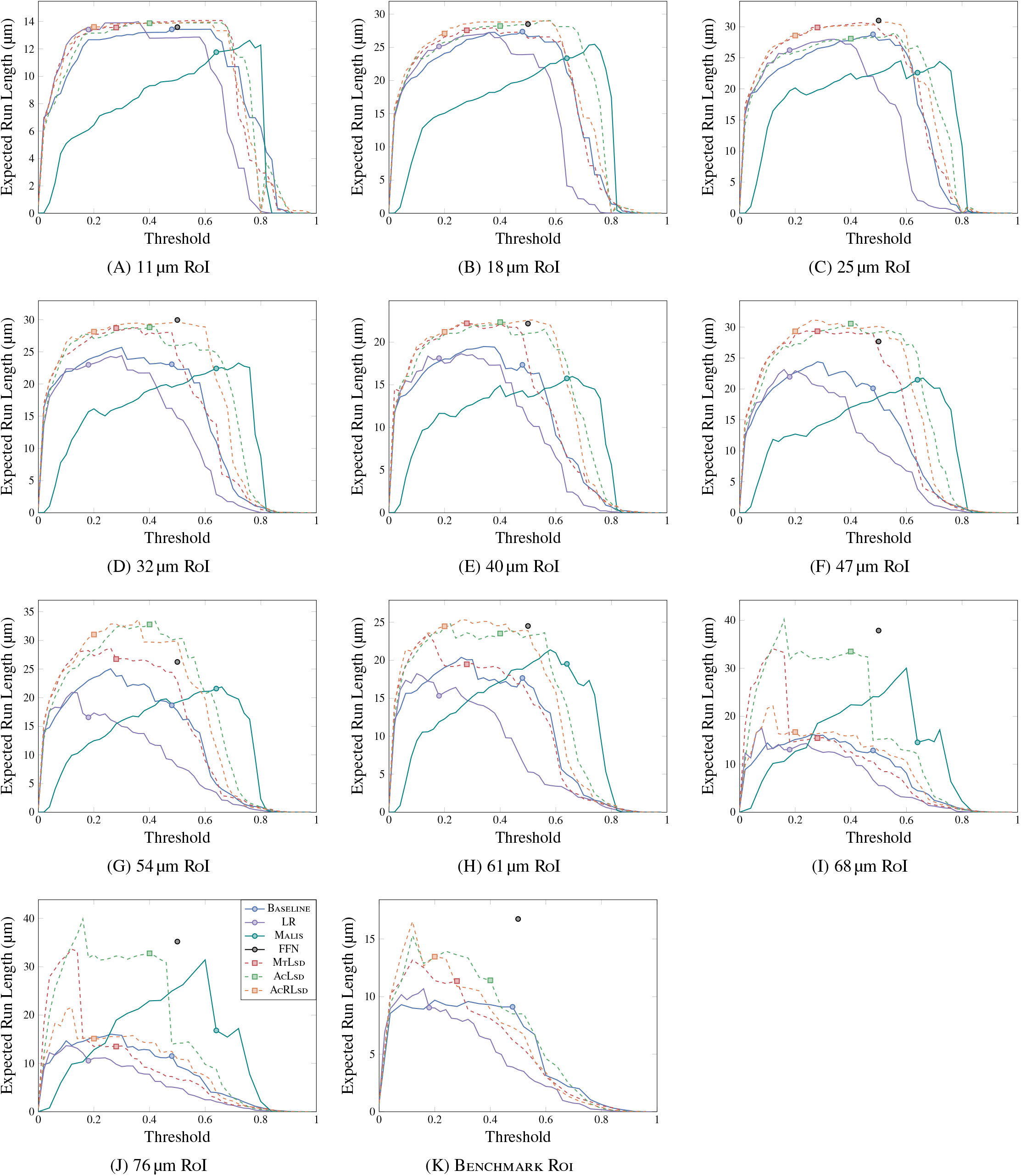
ERL vs threshold across Zebrafinch RoIs. Points correspond to thresholds which maximized ERL on the validation dataset.

**Supplementary Figure 8:**
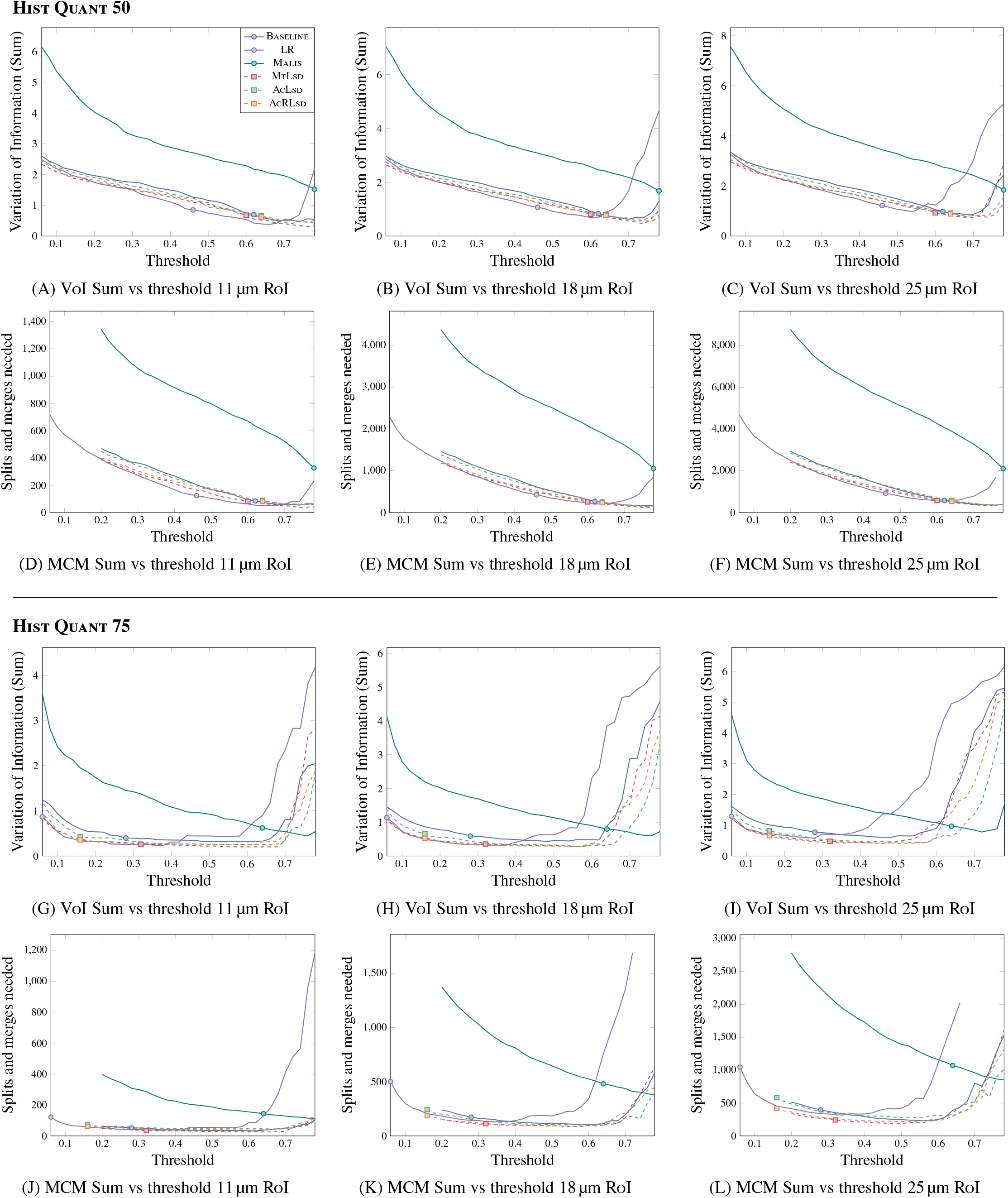
VoI serves as a reasonable proxy for evaluating large volumes. Comparison between VoI and MCM on Zebrafinch on first three RoIs. Similarities are consistent on both Hist Quant 50 (top two rows) and Hist Quant 75 merge functions (bottom two rows). VoI plots are cropped to the threshold range used in the MCM.

**Supplementary Figure 9:**
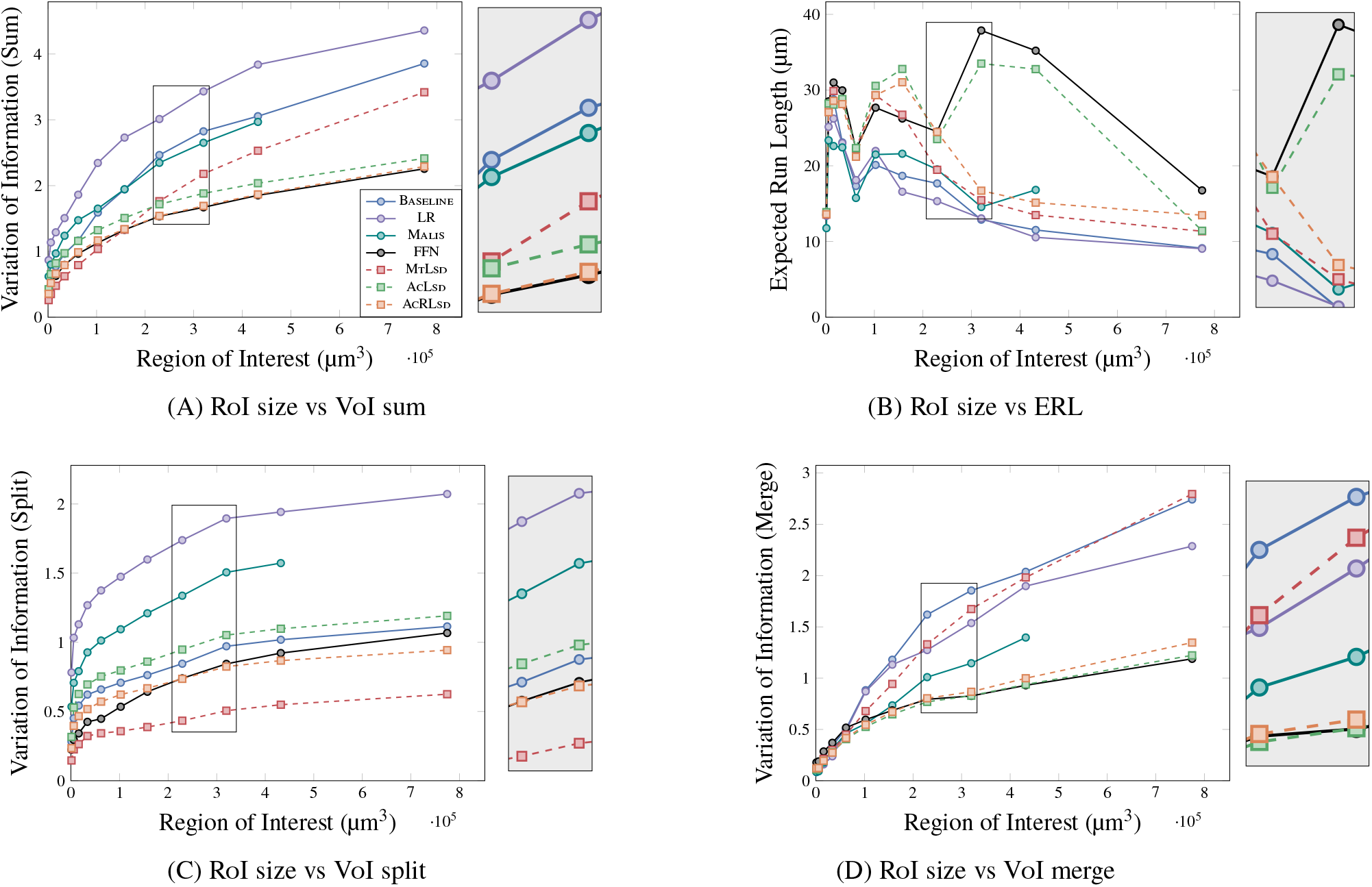
Demonstration of ERL sensitivity. When transitioning from a ∼61µm^3^ to a ∼68µm^3^ RoI, the VoI Sum increases as expected (**A**). This is not the case when considering ERL. All networks, with the exception of FFN and AcLsd, show decreases (**B**). Breaking VoI Sum into false splits (**C**.) and merges (**D**.) shows consistent increases across all networks. However, not all networks reflect this change when considering ERL. This can be best explained by the fact that ERL is more sensitive to different types of merges. In this example, AcRLsd likely merged a small fragment into a larger neuron while AcLsd likely merged two smaller fragments together. While both cases would produce a similar increase in VoI, the ERL in the former is drastically reduced. This is not reflective of the fact that both cases would require a single split to resolve in the context of a contemporary proofreading tool.

**Supplementary Figure 10:**
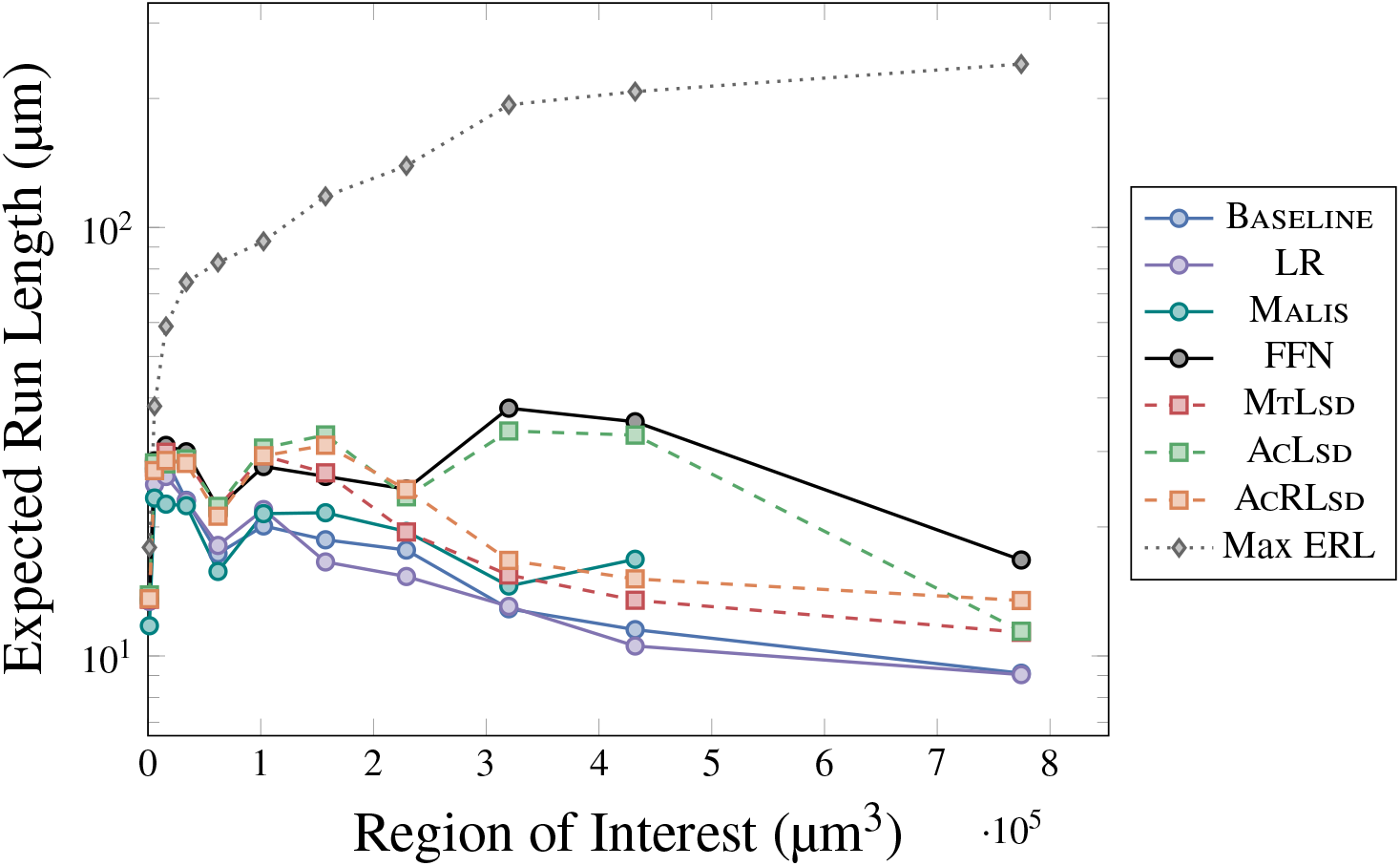
RoI size vs ERL. Grey dotted line shows maximum possible ERL for each RoI. Y-axis is on a log scale

**Supplementary Figure 11:**
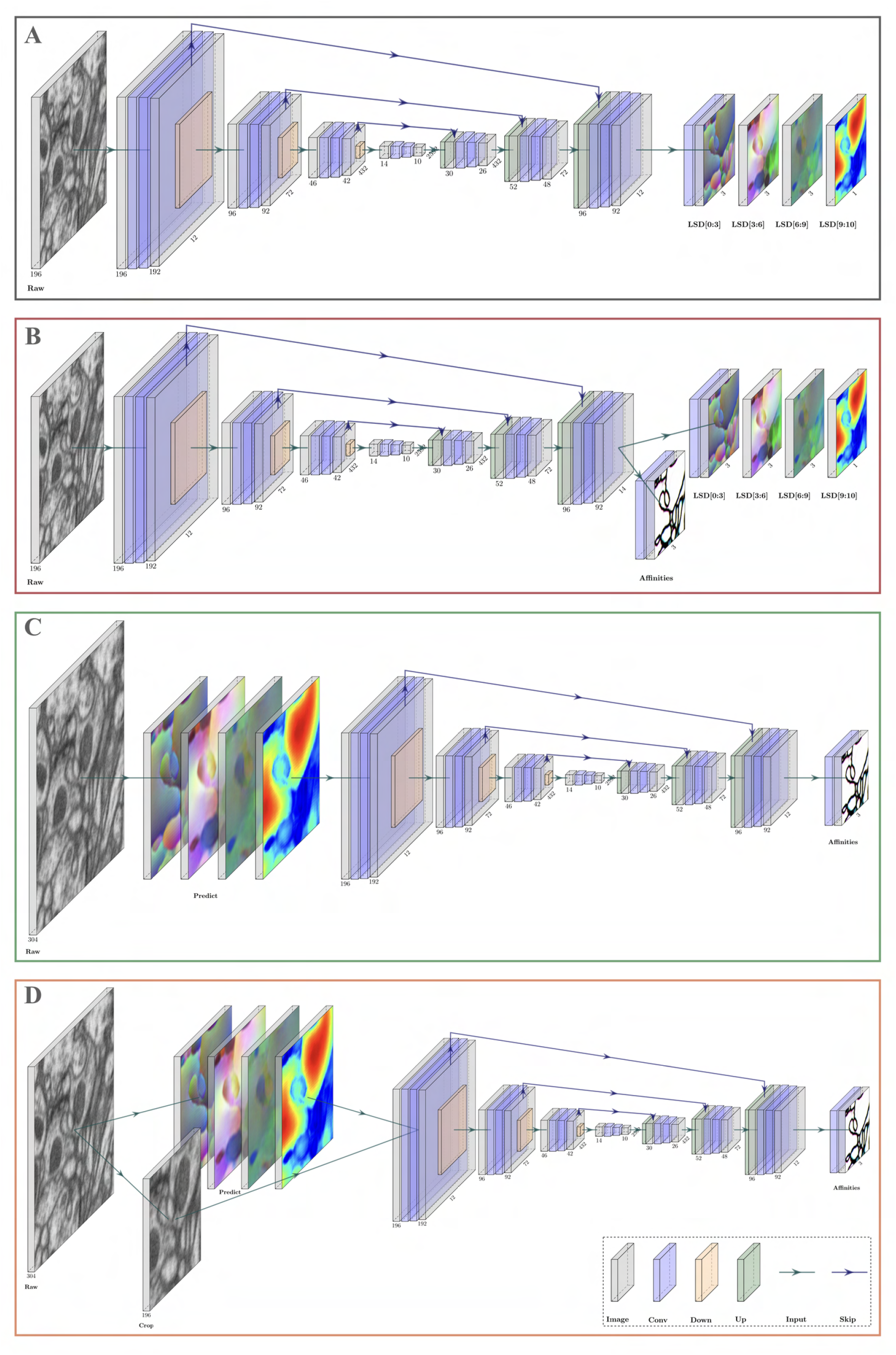
Network architectures used on Fib-SEM datasets (2D representations). All architectures use the U-Net proposed in Funke et al. (2019). Legend on bottom right. **A**. Network to generate the LSDs used in auto-context setups. An extra convolution is used to get to 10 feature maps for the embedding. **B. MtLsd** network - both affinities and LSDs are learnt. Number of output feature maps is increased from 12 to 14 to account for the 13 feature maps needed for the affinities and embedding. **C. AcLsd** network - the output from **A** is used to predict embedding from raw input. The predicted embedding is then passed in to learn affinities. **D. AcRLsd** network - same as **C**, but incorporates cropped raw as input in addition to the embedding.

**Supplementary Figure 12:**
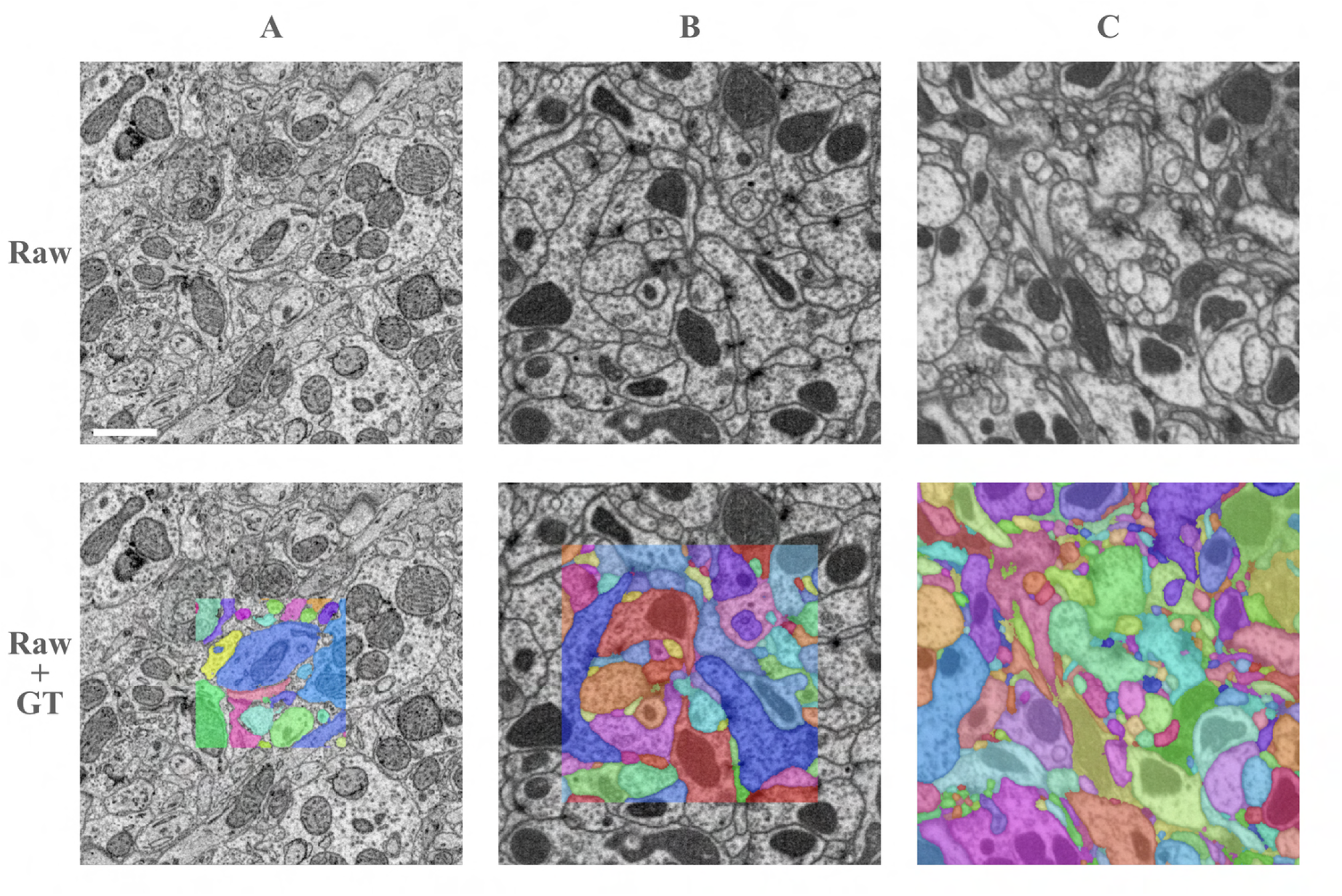
Example training data. **A**. Zebrafinch labels were heavily padded with raw data and some glia were set to zero (scale bar = ∼ 1 µm). **B**. Padding was used to a lesser extent for Hemi-brain volumes. Example taken from Ellipsoid Body. **C**. No padding was used for Fib-25 volumes.

**Supplementary Figure 13:**
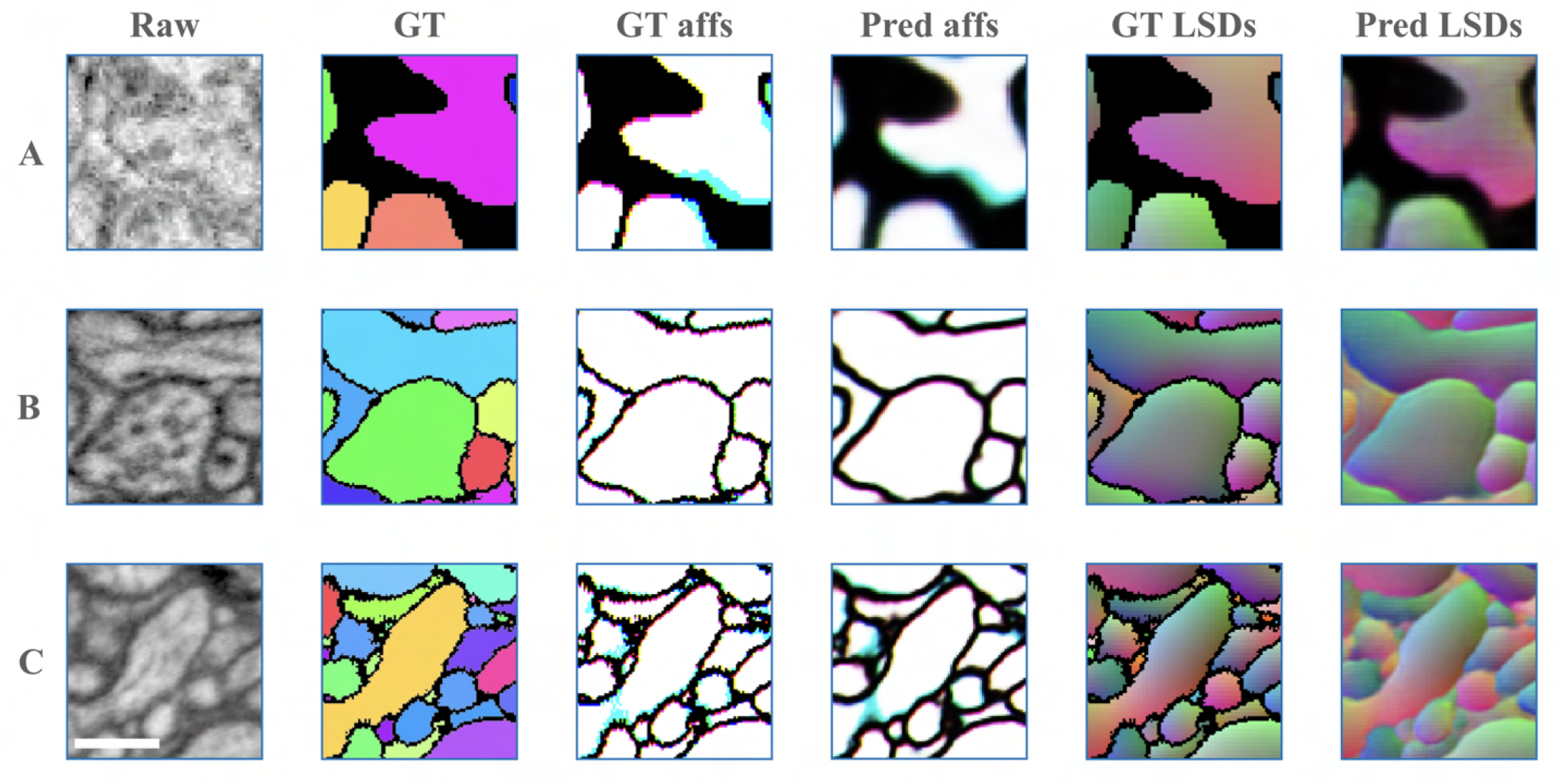
Example training batches. Masked-out regions were factored into the training loss on the Zebrafinch dataset (**A**). Conversely, the Hemi-brain and Fib-25 volumes used no masking during training (**B**,**C**, respectively). Scale bar = ∼300 nm.

**Supplementary Figure 14:**
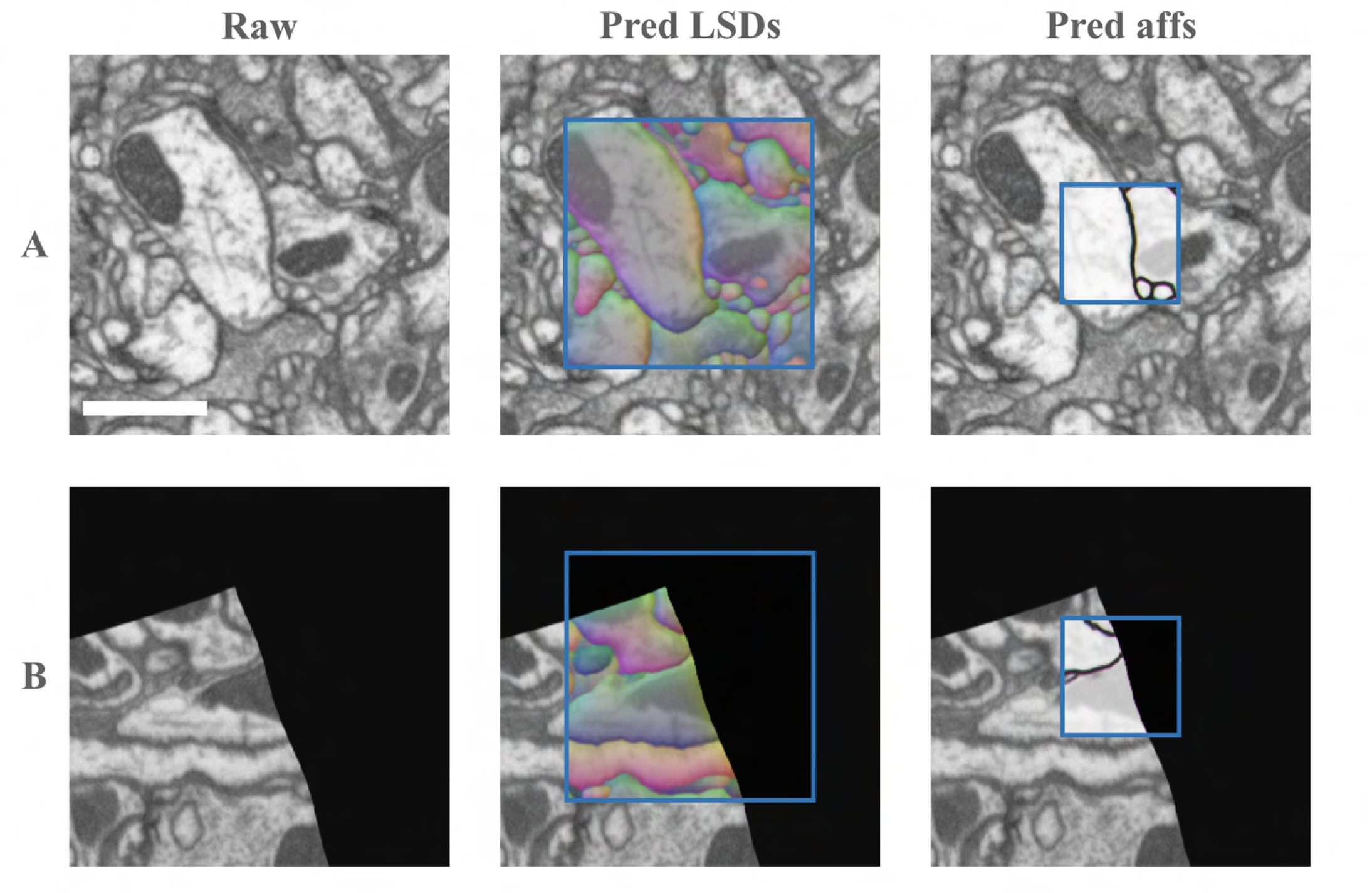
Example auto-context training batches on Fib-25. **A**. Batch in which no boundary masking is needed. The first pass predicts LSDs in an intermediate RoI to provide context for affinity prediction in the second pass. **B**. Batch requires boundary masking. The combination of elastic deformation and zero padding simulates the tissue irregularities seen in the full Fib-25 volume. In these background areas, LSDs and affinities are taught to predict zero. Scale bar = ∼ 1 µm.

**Supplementary Figure 15:**
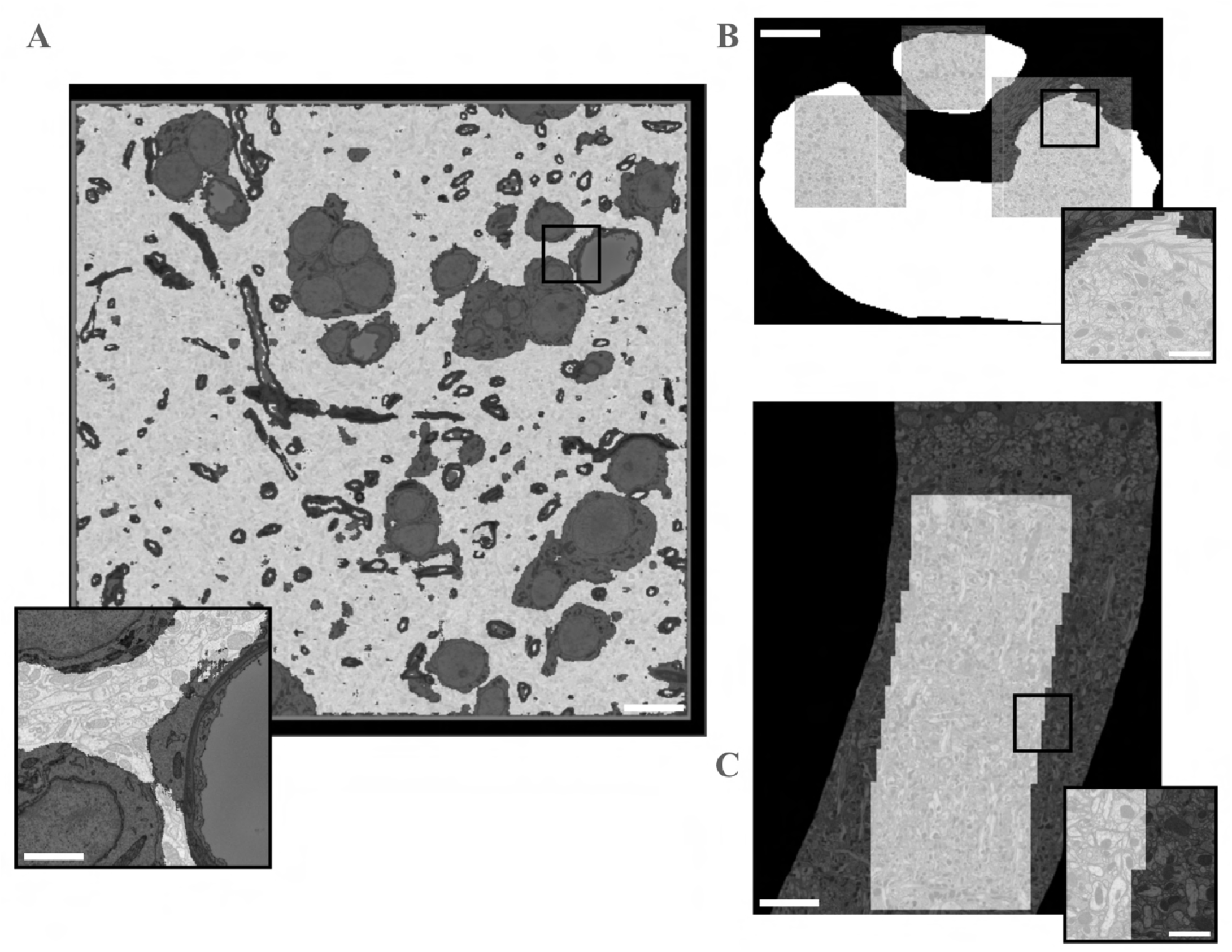
Masks used in study. **A**. Zebrafinch mask removed cell bodies, myelin, blood vessels and background. Scale bar = ∼10 µm. Inset scale bar = ∼3 µm. **B**. Hemi-brain mask restricted volumes to Ellipsoid Body neuropil. Scale bar = ∼ 10 µm. Inset scale bar = ∼2 µm. **C**. Fib-25 used an irregularly shaped tissue mask, mostly limited to neuropil. Scale bar = ∼8 µm. Inset scale bar = ∼1 µm.

**Supplementary Figure 16:**
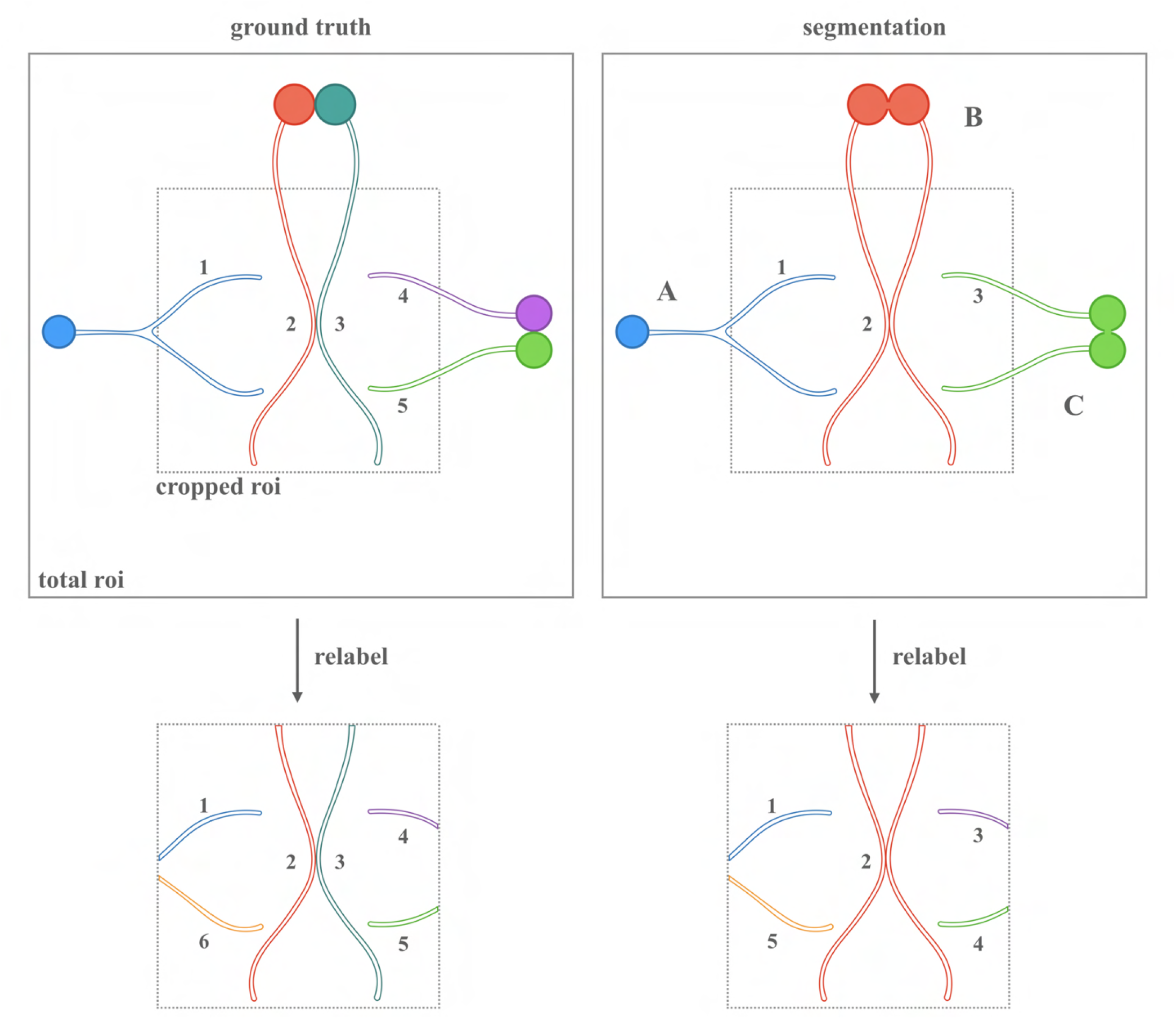
Potential side effects of cropping data. Ground truth (left) shows 5 correctly labeled neurons. Example Segmentation (right) shows 3 labeled neurons as a result of false merges. Bottom squares show results from relabeling connected components inside the cropped RoI. **A**. Correctly segmented neuron in total RoI would be counted as a false merge inside cropped RoI. This is fixed by relabelling connected components and the merge/split scores are unaffected. **B**. A falsely merged neuron inside the cropped RoI is caused by a false merge outside of the cropped RoI and should not be counted. Relabelling doesnt resolve the touching boundaries and the merge score is subsequently overestimated. **C**. Incorrectly segmented neuron in total RoI is counted as correct in cropped RoI after relabelling. Merge score is underestimated.

**Supplementary Figure 17:**
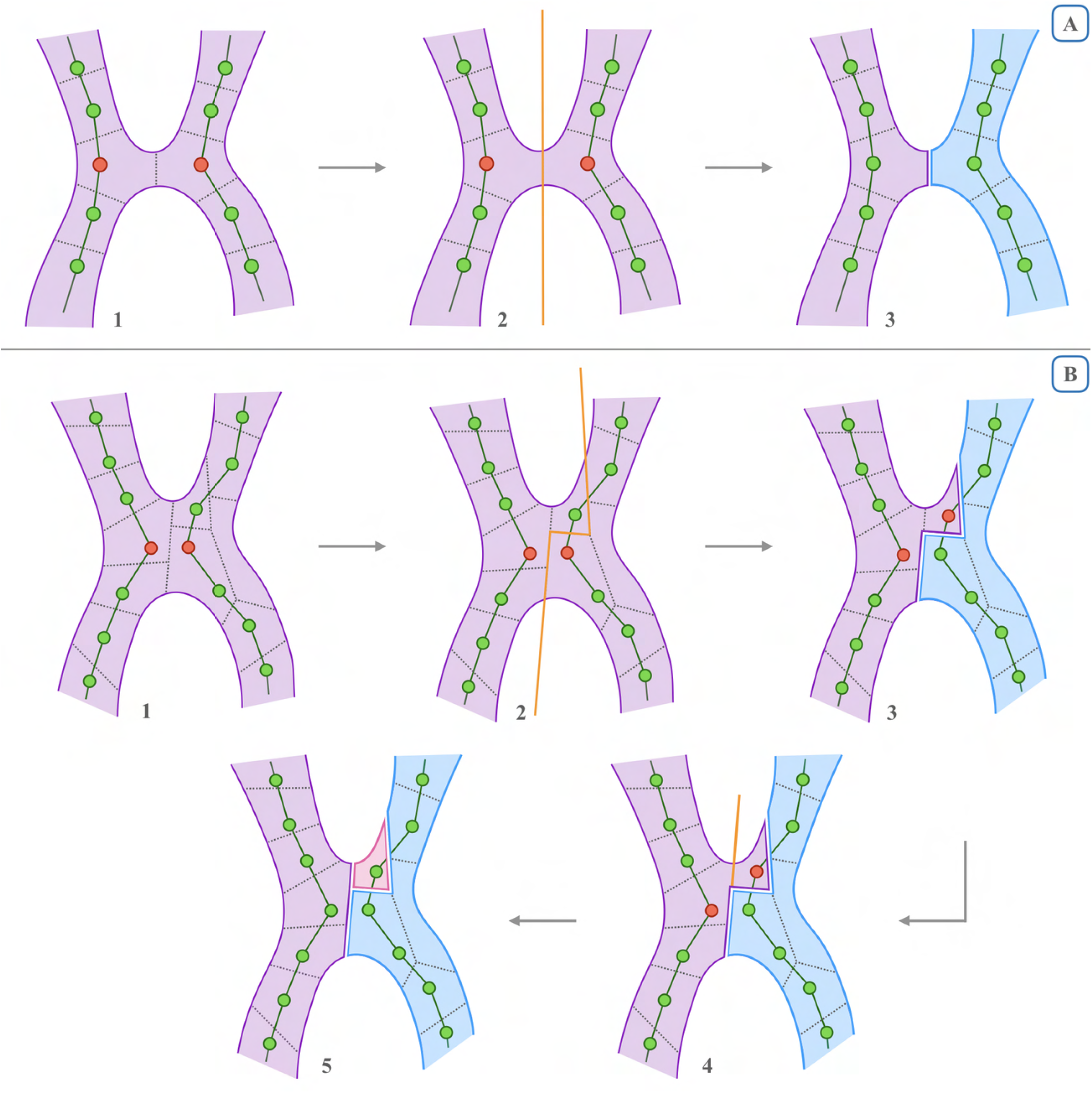
Overview of the proposed Min-cut Metric (MCM). **A**. Simple case. Two ground-truth skeletons are contained inside an erroneously merged segment. Dashed lines represent supervoxel boundaries and the closest skeleton nodes need to be split to resolve the merge (1). A min-cut is performed (2), resulting in a new segment (3). **B**. Complex case. Two skeletons are contained in a falsely merged segment as before (1), but the supervoxels are more fragmented. A min-cut is performed (2), resulting in a new segment (3). However, two nodes contained within the original segment need to be split. A second min-cut is performed (4), which produces another segment (5). This results in an additional split error caused by the original cut.

**Supplementary Figure 18:**
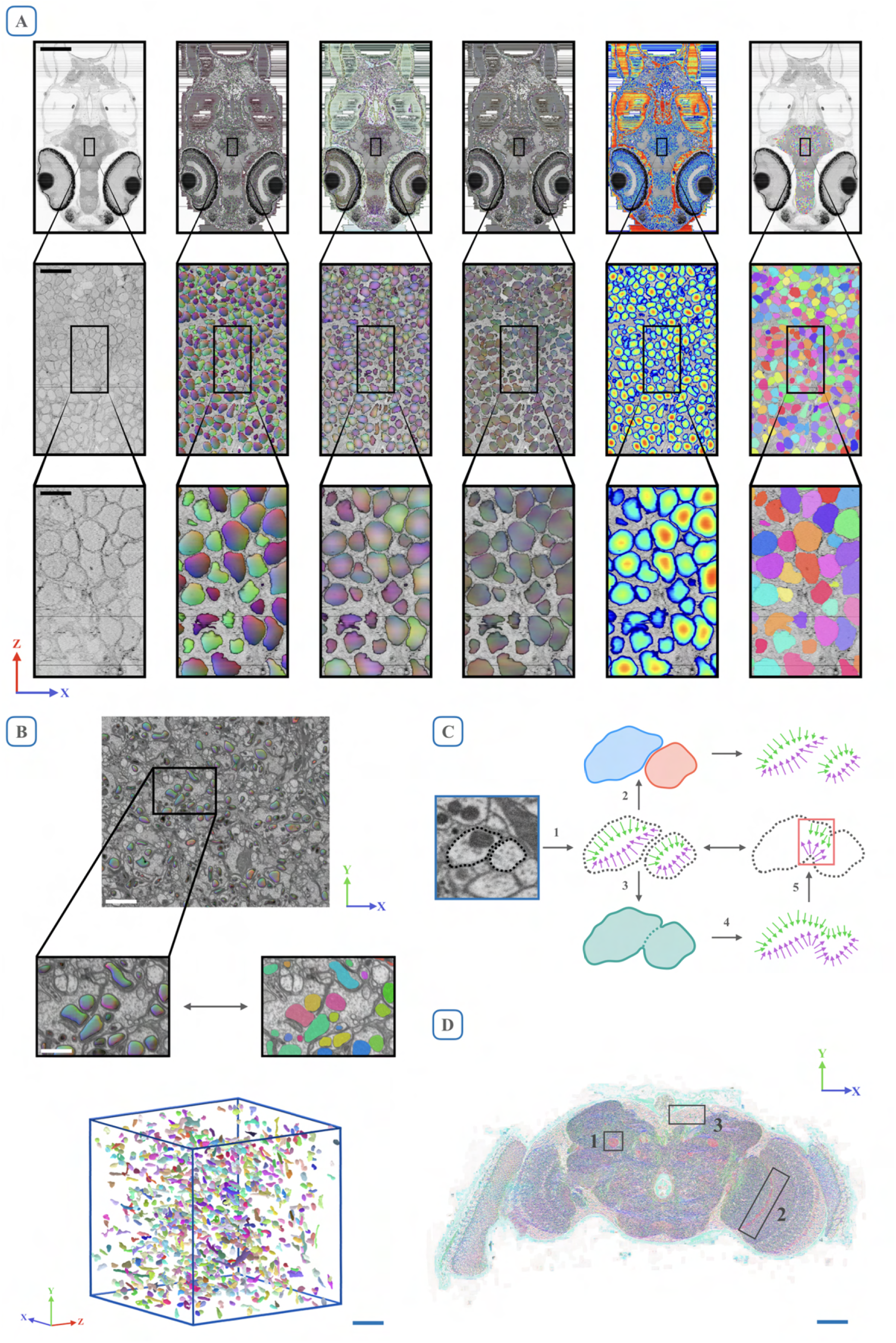
Other potential uses of LSDs. **A**. Nuclei segmentation on full zebrafish brain (Hildebrand et al., 2017). Columns from left to right: raw, LSD offset vectors, LSD direction vectors (covariance), LSD direction vectors (Pearson’s), size, resulting segmentation. Scale bars from top to bottom: ∼ 150µm, 20µm, 5µm. **B**. Mitochondria segmentation on cropout from Fib-25. Inset shows LSD predictions and corresponding segmentation. Bottom image shows 3D reconstructions of a random sample (n=1000) in predicted RoI. Scale bars from top to bottom: ∼ 3µm, 750nm, 4µm. **C**. Error mapping. Example predicted LSDs between two neurons (1). If the resulting segmentation is correct (2), segmentation LSDs do not differ from predicted LSDs. If the resulting segmentation is incorrect (3), segmentation LSDs (4) might differ from the predicted LSDs. The difference (5) could expose errors in a segmentation. **D**. Predicted direction vectors (covariance) on single section of full adult fly brain. Mushroom body pedunculi (1), optic chiasm (2), cell rind (3) highlight directionality. Scale bar = ∼ 150µm.

## Footnotes

1 https://github.com/funkelab/lsd

2 See https://www.ilastik.org/documentation/autocontext/autocontext for a popular example of this strategy.

3 Distinct from shape descriptors in Maitin-Shepard et al. (2016).

4 http://funkey.science/gunpowder

5 https://www.tensorflow.org/

6 Kindly provided by the authors of Januszewski et al. (2018)

7 Kindly provided by the Janelia FlyEM project team (https://janelia.org/project-team/flyem)

8 The authors of Januszewski et al. (2018) report a VoI split of 0.8837. While we were able to replicate the reported VoI merge score of 0.053, we found VoI split to be 1.003.

9 https://github.com/funkelab/lsd

10 Also shown by the authors of Januszewski et al. (2018) in supplementary tables 4 and 5.

11 https://cremi.org

12 http://brainiac2.mit.edu/SNEMI3D

13 Also referred to as “cleaving”.

14 MtLsd network had 14 output feature maps to account for 13 final feature maps from the affinities (3) and LSDs (10)

15 https://github.com/funkey/waterz

16 https://github.com/funkelab/funlib.segment

17 https://github.com/funkelab/funlib.evaluate

18 https://github.com/seung-lab/cloud-volume

19 Ellipsoid Body: 2, Protocerebral Bridge: 2, Fan-Shaped Body: 2, Lobula Plate: 1, Lateral Horn: 1

20 Mask contains some background and cell bodies

21 https://github.com/janelia-flyem/dvid

22 https://github.com/janelia-flyem/neuclease

23 One block is defined as the largest output volume that can be predicted by a network in one pass on the respective GPU it was evaluated on.

24 https://www.tensorflow.org/guide/profiler

## Notes

### Competing Interest Statement

The authors have declared no competing interest.

### Summary of Updates

Added experiments to test LSD efficacy on mouse brain serial section EM and non-EM (plant epithelial cell) data. Added ablation experiment to gauge importance of combinations of the LSD components. Reformulated equations, added text, fixed various figure axes, added citations.

